# Wildebeest migration in East Africa: Status, threats and conservation measures

**DOI:** 10.1101/546747

**Authors:** Fortunata U. Msoffe, Joseph O. Ogutu, Mohammed Y. Said, Shem C. Kifugo, Jan de Leeuw, Paul Van Gardingen, Robin S. Reid, JA Stabach, Randall B. Boone

## Abstract

Migration of ungulates is under pressure worldwide from range contraction, habitat loss and degradation, anthropogenic barriers and poaching. Here, we synthesize and compare the extent of historical migrations of the white-bearded wildebeest (*Connochaetes taurinus*) to their contemporary status, in five premier East African ecosystems, namely the Serengeti-Mara, Masai Mara, Athi-Kaputiei, Amboseli and Tarangire-Manyara. The current status, threats to migration, migratory ranges and routes for wildebeest were characterized using colonial-era maps, literature reviews, GIS and aerial survey databases, GPS collared animals and interviews with long-term researchers. Interference with wildebeest migratory routes and dispersal ranges has stopped or severely threatens continuation of the historical migration patterns in all but the Serengeti-Mara ecosystem where the threat level is relatively lower. Wildebeest migration has collapsed in Athi-Kaputiei ecosystem and is facing enormous pressures from land subdivision, settlements and fences in Amboseli and Mara ecosystems and from cultivation in Tarangire-Manyara ecosystem. Land use change, primarily expansion in agriculture, roads, settlements and fencing, increasingly restrict migratory wildebeest from accessing traditional grazing resources in unprotected lands. Privatization of land tenure in group ranches in Kenya and settlement policy (villagization) in Tanzania have accelerated land subdivision, fencing and growth in permanent settlements, leading to loss of key wildebeest habitats including their migratory routes and wet season calving and feeding grounds. These processes, coupled with increasing human population pressures and climatic variability, are exerting tremendous pressures on wildebeest migrations. Urgent conservation interventions are necessary to conserve and protect the critical wildebeest habitats and migration routes in East Africa.

## Introduction

Large mammal migrations are among the most-awe inspiring of all migrations [1]. These migrations, the seasonal and round-trip movement of large herbivores between discrete areas, are under increasing pressures worldwide. Globally, migrations of 6 out of 24 species of ungulates are either already extinct or their status is unknown [2]. Of the remaining ungulate mass migrations, most occur in six locations in Africa, including the white-bearded wildebeest (*Connochaetes taurinus* Burchell, 1823) migration in the Serengeti-Mara ecosystem of Kenya and Tanzania [2]. Range restriction and alteration, degradation and loss of habitat due to agriculture, poaching and barriers that block migration, such as fences, roads, railroads, pipelines and settlements have progressively disrupted historical migratory routes and decimated or driven rapid population declines of many of the once spectacular migratory herds over the 20^th^ century [1,3–5].

Because migration enables populations to grow to large abundances, its disruption leads to restricted ranges and consequent population declines [1,6,7]. The preservation of the phenomenon of migration requires conservation of both the migratory species and the habitats along their routes. It also requires a sound understanding of the factors and processes underlying the degradation and loss of migratory routes and declines of populations to devise effective strategies for protecting migratory routes, habitats and populations [1]. Although causes of ungulate migrations are not yet fully understood [8], the temporal regularity of migrations suggests that they are a response to seasonal fluctuations in spatial patterns of resource availability and quality [9,10]. Thus, in the Serengeti-Mara ecosystem, rainfall through its effect on food supply and salinity of drinking surface water has been suggested as a trigger for the northward migration [11] whereas high nutrient availability on the short grass plains is thought to attract lactating female wildebeest southwards [12]. This migration results in the movement of wildebeest from the open, highly nutritive grasslands with low biomass in the wet season, to wooded grasslands with high biomass of lesser nutritive quality during the dry season [13].

We focus on populations of wildebeest sub-species in East Africa because they (1) are taxonomically closely related, (2) represent some of the most important remaining large mammal migrations on earth, (3) share similar conservation problems, (4) all have ranges within and outside protected areas, and (5) have a range of current and potential pathways to protection. The threats facing wildebeest migrations involve the interplay of multiple factors and processes [1–3,14]. In the Masailand of Kenya and Tanzania, ungulate population declines, particularly of wildebeest, are linked to habitat loss due to land use change or habitat degradation caused mainly by expansion of cultivation [9,15,16]. Illegal hunting might, however, have contributed more to dwindling populations of migratory ungulates in some areas, including the Tarangire-Manyara ecosystem of Tanzania [17–19]. Wildebeest also cause problems for livestock, including competition for forage and transfer of the deadly malignant catarrhal fever virus from wildebeest calves to cattle [20,21]. The type and intensity of these factors and processes vary among migratory species and across their meta-populations or ecosystems. Effective wildlife conservation and protection thus requires clear prioritization of the factors leading to population declines both in the short-and long- term. Integral to this process is reviewing the history, status, trends and threats facing populations of particular migratory species across a range of ecosystems along the entirety of their migratory routes to extract general insights into the threats they face as a basis for developing approaches likely to succeed in conserving their populations and migrations.

We describe and compare the extent of historical migrations of the western (*C.t. mearnsi*) and eastern (*C.t. albojubatus*) white-bearded wildebeest with the current status of these migrations and migratory routes in five ecosystems of East Africa. We evaluate long-term wildebeest population trends, putative drivers of change and their impacts on the critical habitat and migratory ranges of wildebeest in each of the five ecosystems. We suggest potential strategies for conserving these migrations, some of which rank among the Earth’s most spectacular remaining terrestrial migrations (Fig 1). Lastly, we evaluate causes of wildebeest population declines and range contraction, including human population expansion, land-use change, poaching, land uses incompatible with wildlife conservation, deficiencies in existing wildlife policies, institutions and markets in Kenya and Tanzania and suggest conservation strategies to alleviate the population declines.

**Fig 1.**
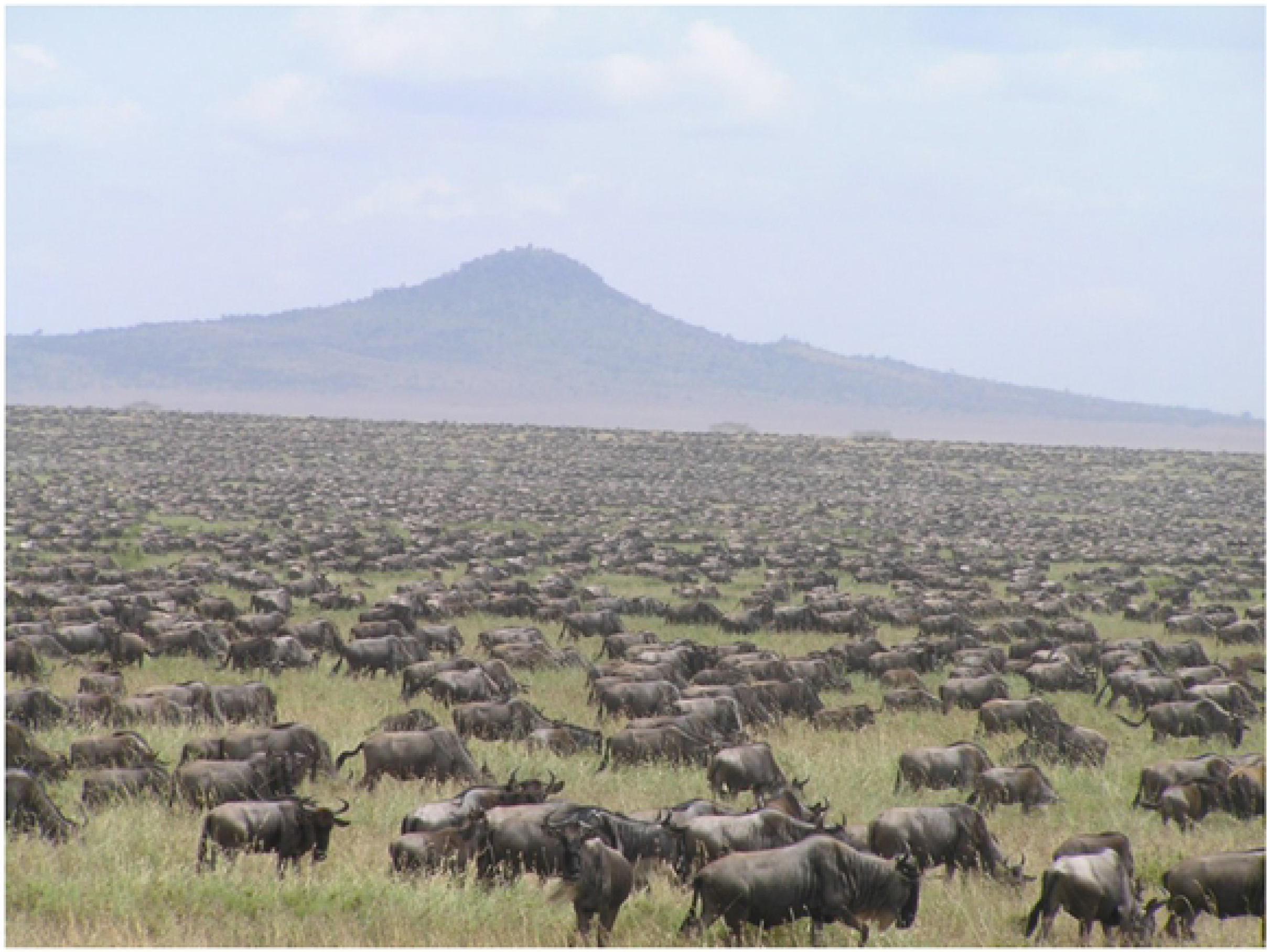
Wildebeest migration in the greater Serengeti-Mara ecosystem in East Africa. (Photo credit: ARE Sinclair)

## Materials and methods

### Study Area

This study covers the five ecosystems in East Africa with migratory wildebeest populations (Fig. 2). These include the Serengeti-Mara, Loita Plains, Athi-Kaputiei Plains, Amboseli Basin, and Tarangire-Manyara ecosystems. Across these five ecosystems, we focus on eight populations of either the western (Serengeti-Mara, Ngorongoro, Loita Plains, Narok County) or the eastern (Athi-Kaputiei, Machakos County, Amboseli, West Kajiado, Tarangire-Manyara) subspecies of the white bearded wildebeest [22]. We consider three (Ngorongoro, Narok County and Machakos) of the eight populations only superficially because they are part of at least one of the other populations considered in detail. We do not consider small, resident wildebeest populations occupying the western corridor in Serengeti and the Loliondo Game Controlled Area (LGCA) in north-eastern Tanzania [14].

**Fig 2.**
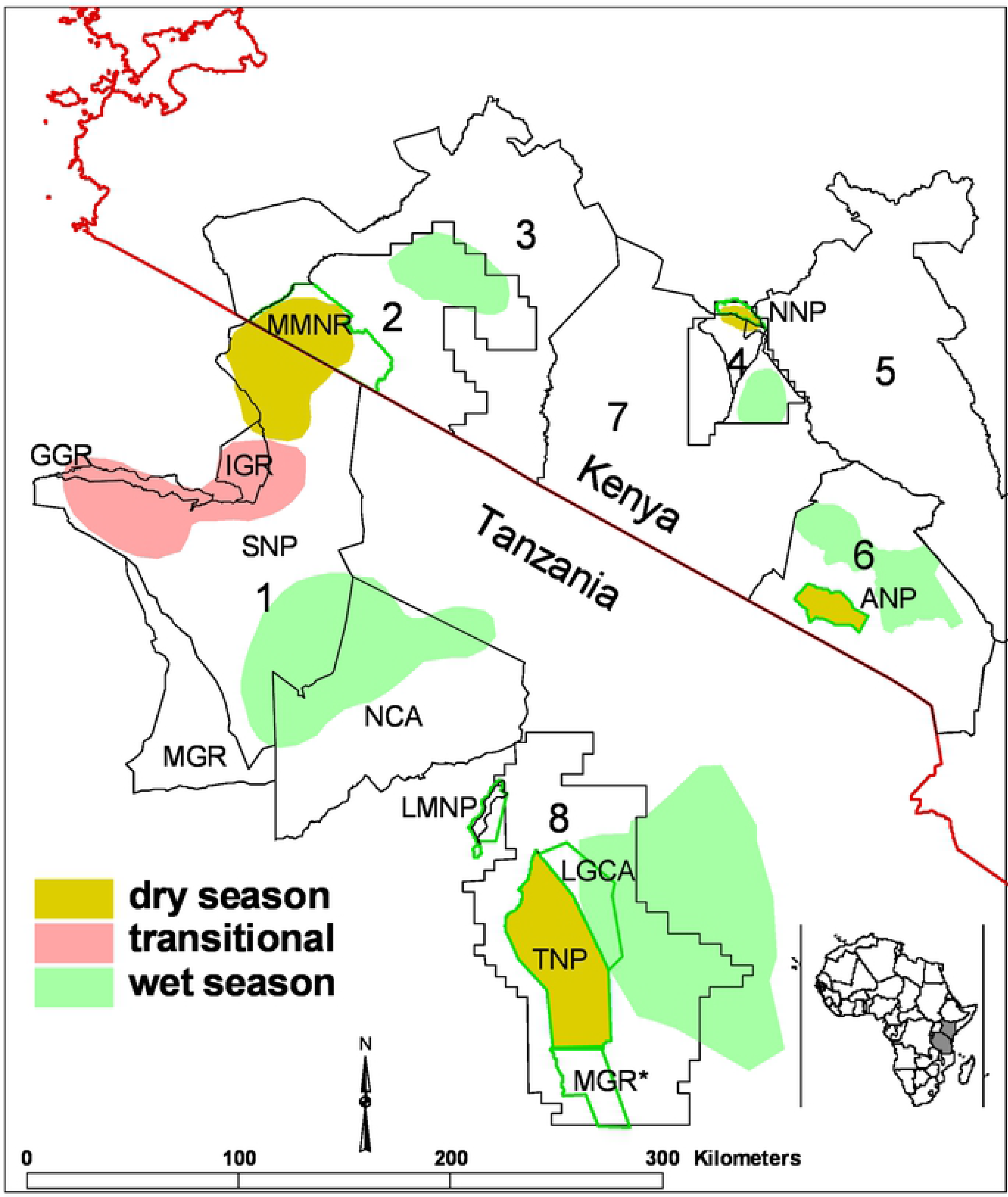
Map showing the general extent of Masailand in Kenya and Tanzania and the five study ecosystems with eight populations: 1 = Serengeti Ecosystem, 2 = Masai Mara Ecosystem, 3 = Narok County, 4 = Athi-Kaputiei Ecosystem, 5 = Machakos County, 6 = Greater Amboseli Ecosystem, 7 = West Kajiado and 8 = Tarangire-Manyara Ecosystem populations. Notes: NCA = Ngorongoro Conservation Area, MGR = Maswa Game Reserve, SNP =Serengeti National Park, IGR = Ikorongo Game Reserve, GGR = Grumeti Game Reserve, MMNR = Masai Mara National Reserve, NNP = Nairobi National Park, ANP = Amboseli National Park, LMNP = Lake Manyara National Park, LGCA = Lokisale Game Controlled Area, TNP = Tarangire National Park and MGR* = Mkungunero Game Reserve. Use of each seasonal area by the study populations is described in the text.

The Serengeti-Mara Ecosystem covers about 40,000 km^2^ in Tanzania and Kenya [14,23]. The ecosystem encompasses the Serengeti National Park, Ngorongoro Conservation Area, Maswa, Grumeti, Ikorongo and Kijereshi Game Reserves, Loliondo Game Controlled Area, Ikona and Makao Wildlife Management Areas in Tanzania and the Masai Mara National Reserve and adjoining wildlife conservancies and pastoral ranches in Kenya.

The Ngorongoro Conservation Area (NCA, 8,292 km^2^) is part of the Greater Serengeti-Mara ecosystem. It includes the Ngorongoro Crater (310 km^2^) and is bordered to the north by the Loliondo Game Controlled Area (4000 km^2^). Lake Natron Game Controlled Area (LNGCA, 3000 km^2^) borders the LGCA to the southeast and the NCA to the northeast (Fig. 2).

The Narok County (17,814 km^2^) encompasses the Loita Plains and the Masai Mara Ecosystem in Kenya. The Athi-Kaputiei ecosystem (2,200 km^2^) covers the Nairobi National Park (117 km^2^) and the adjacent Athi-Kaputiei Plains in Kenya. Machakos County (14,225 km^2^) is contiguous with the Athi-Kaputiei ecosystem. The Greater Amboseli ecosystem of Kenya (7730.32 km^2^) covers the Amboseli National Park (392 km²) and surrounding dispersal areas on pastoral rangelands, covering some 3,000 km^2^ [24–26]. Western Kajiado (11388.54 km^2^) is bounded by the Greater Amboseli Ecosystem to the East. Both ecosystems are found in Kajiado County of Kenya.

The Tarangire-Manyara ecosystem of Tanzania covers the Tarangire (2,850 km^2^) and Lake Manyara (649 km^2^) National Parks and Manyara Ranch (177 km^2^), a private conservancy that supports livestock rearing, wildlife conservation and tourism. This ecosystem is adjoined by rangelands managed primarily for cultivation, livestock grazing, legal game hunting, and tourism on community land designated as Open Areas, Game Controlled Areas or Wildlife Management Areas [15]. These include the Simanjiro Plains, the Mkungunero Game Reserve (800 km^2^) and Lolkisale Game Controlled Area (1500 km^2^). Altogether, the range for the migratory wildebeest covers about 35,000 km^2^ [17,27–30].

Human population growth drives sedentarization, expansion of settlements, fences and other land use developments in the study ecosystems [4,5]. These changes promote land use intensification and illegal livestock incursions into protected areas to the detriment of migratory wildebeest [31,32]. In Kenya, human population size increased in Narok County by 673% from 110,100 in 1962 to 850,920 in 2009; in Kajiado County by 905% from 68,400 in 1962 to 687,312 in 2009 and in Machakos County by 247% from 571,600 in 1962 to 1,983,111 in 2009 [33]. Similarly, in Tanzania human population size increased in the Serengeti District by 11.6% from 249,420 in 2012 to 282,080 in 2017 and in Monduli and Simanjiro Districts, containing the Tarangire-Manyara Ecosystem, by 13.5% from 460,775 people in 2012 to 532,939 in 2017 (www.nbs.go.tz).

Across the Serengeti-Mara ecosystem, Narok County, Masai Mara ecosystem and the Loita Plains, rainfall is markedly bimodal and increases steeply along a southeast–northwest gradient, from east to west, south to north and over time [34]. Notably, rainfall increases from 500 mm on the Serengeti Plains to the Southeast to 1400 mm to the north-west of Masai Mara National Reserve. Across the Kajiado County in which the Amboseli, Athi-Kaputiei and Western Kajiado Ecosystems are found, rainfall is low, bimodal and highly variable, and total annual rainfall averages 685 mm (range 327-1576 mm). The short rains fall from November to December (30.97 ± 27.85% of the annual total) and the long rains from March to May (47.5 ± 15.06% of the annual total). The dry season rains fall during June-September. Rainfall is markedly variable in space and increases with elevation such that it averages 300 mm/yr in the low-lying Amboseli basin and rises to 1250 mm/yr on the slopes of Mt. Kilimanjaro and Chyulu Hills in the southeast of the County to 800 mm in Nairobi National Park and 971 mm at Ngong hills in the northwest of the County [26]. Rainfall increases from under 500 mm in the extreme southeast of the Athi-Kaputiei Plains to over 800 mm in northern Nairobi Park [35]. In the Tarangire-Manyara ecosystem, rainfall is bimodal and averages 650 mm per annum. The short rains span from October to December and the long rains from March to May. The rains are unreliable and frequently fail, especially the short rains [15]. Land use patterns in the study ecosystems are described comprehensively elsewhere [15,26,34–36].

### Historical wildebeest migrations in East Africa

Information on the migratory wildebeest range, routes and status was compiled from literature reviews, colonial-era records, maps, GIS databases, Global Positioning System (GPS) collared wildebeest and interviews with local residents and researchers knowledgeable about the study ecosystems. We reviewed historical records to provide a context for assessing changes in wildebeest migrations in East Africa.

### Mapping contemporary wildebeest migratory routes and ranges

To obtain information on contemporary wildebeest movements, we placed GPS collars on 15 wildebeest in the Loita Plains in the Mara Ecosystem in May 2010, 12 in the Athi-Kaputiei Plains and 9 in the Amboseli Basin in October 2010. The collars were programmed to collect the position of each wildebeest 16 times each day (every hour from 6:00 AM to 6:00 PM and every three hours from 6:00 PM to 6:00 AM) for a 2-year study period. Data are available on Movebank (www.movebank.org)

In the Tarangire-Manyara ecosystem, OIKOS and Tanzania National Parks [37] tracked movements of radio collared wildebeest and zebra and GPS collared elephants (*Loxodonta africana*) during 1995-2002 to establish if they still used the main migratory routes identified earlier [38]. OIKOS also established the presence or absence of migratory routes and assessed wildlife species abundance in the ecosystem during 1995-2002 by interviewing local communities, hunting operators, employees and residents and conducting multiple aerial reconnaissance and systematic reconnaissance flights. Several studies later mapped and analysed land use changes along the migratory corridors [36,39]. We did additional unstructured interviews on the status of the migration routes in the ecosystem during 2006-2007. Our interviews targeted long-term local residents and researchers and were carried out during ground truthing work for imagery analysis on historical land use and cover changes in the ecosystem from 1984 through 2000 to 2006-2007. Local Masai elders who knew the history of the ecosystem well helped with the ground truthing and interviewing local residents about land use and cover changes. The field data form used for our interviews is provided in Table S1.

### Wildebeest population trends

Wildebeest population estimates were compiled from aerial surveys conducted in Kenya by the Directorate of Resource Surveys and Remote Sensing (DRSRS) and in Tanzania by the Tanzania Wildlife Research Institute (TAWIRI), Tanzanian Wildlife Conservation Monitoring Unit (TWCM) and Frankfurt Zoological Society (FZS). The methods used in the aerial surveys and for estimating population size are described in detail elsewhere [33,40–42]. Aerial surveys began in the Athi-Kaputiei ecosystem in 1949 [43], in the Serengeti-Mara ecosystem in 1957 [44–46]), in the Tarangire-Manyara ecosystem in 1964 [27] and in Amboseli in 1973 [47].

### Distribution of cultivation and fences

Data on the distribution of agriculture were obtained from the FAO Africover project 2000 [48]. The project mapped land cover for the year 2000 for the whole of East Africa from Landsat images (30 m resolution) and updated the Kenya map in 2008. The map category ‘agriculture’ was extracted from the Africover data set and clipped according to the study area boundary. In the Athi-Kaputiei ecosystem fences were mapped in 2004 and 2009 by the International Livestock Research Institute (ILRI) and African Wildlife Foundation (AWF) in collaboration with the local communities and local NGO’s using hand-held (GPS, with scientific, technical and logistical support provided by ILRI [4,5,35]. Fences, settlements, roads and other infrastructures were similarly mapped with hand held GPS in Amboseli in 2004-2006 [49,50] and in Masai Mara in 1999, 2002 and 2015 [51–54]. A few fences also exist in the ecosystem in Tanzania.

### Wildlife conservation initiatives and gaps in policies, institutions and markets in Kenya and Tanzania

We reviewed official records on contemporary wildlife conservation initiatives and identified important gaps in wildlife policies, institutions and markets in Kenya and Tanzania.

### Statistical Analysis

Estimates of wildebeest population size for each ecosystem were obtained using Jolly’s method II for transects of unequal lengths [55] and related to the year of survey using negative binomial regression models with linear and quadratic polynomial terms and serial autocorrelation in the counts accounted for using the first-order autoregressive model. Selection between the linear and quadratic models was based on the Akaike Information Criterion [56]. The models were fitted using the SAS GLIMMIX procedure [57]. Temporal trends in wildebeest population size were modeled using a semiparametric generalized linear mixed model with a negative binomial error distribution and a log link function in SAS GLIMMIX procedure [33]. The percentage change in population size between the start and end dates of the surveys was estimated for each ecosystem. For some ecosystems predicted population size for two to three consecutive surveys were averaged and used to compute the averages to minimize the effect of stochastic noise due to small sample size or areal coverage.

## Results

### Historic and contemporary migratory routes

The seasonal migration of the white-bearded wildebeest and zebra (*Equus quagga burchelli*) from the Serengeti Plains in Tanzania to Masai Mara in Kenya [14,46] is called the southern migration. It differs from the northern migration that involves seasonal movements of wildebeest, zebra and Thomson’s gazelle (*Gazella thomsoni*) between the Loita Plains and Masai Mara National Reserve within the Narok County of Kenya [9,10,16,58]. The wildebeest involved in the northern migration form the bulk of the Narok County population.

The migration ranges, routes and population trends of the Serengeti-Mara wildebeest have been extensively studied (e.g., [59,60], Table S2). After rinderpest killed about 95% of this population between 1890 and 1892, it remained low till 1962. The population increased from 263,362 in 1961 to 483,292 in 1967 following veterinary removal of rinderpest in wildebeest in 1962 and again from 1967 to 1.4 million in 1977, coincident with an increase in the dry season rainfall [61,62].

The migration pattern of the Serengeti-Mara wildebeest in the 1940s and 1950s was different from what it is today. Then, wildebeest migrated periodically between the Kenya’s Loita Plains and Tanzania [63,64]. Heavy harvesting of wildebeest (plus zebra and other species) in Narok County during World War II to provide meat for labour and prisoners of war and reduce competition with Masai livestock until 1947 [64 reduced their population. Even so, wildebeest population on the Loita Plains numbered about 50,000-100,000 individuals prior to 1947 [45]. Thereafter, the population further declined drastically because the Loita Plains were opened up to uncontrolled commercial meat-hunters for a short period after World War II [67]. Subsequently, the Mara population (including the Loita Plains) numbered about 15000 by 1958 [45] and 17,817 by 1961 [65].

The Mara wildebeest [45,46,58,66] were formerly more numerous and their distribution extended far beyond their contemporary range in the Mara region of Narok County. But major land use and cover changes have progressively degraded and reduced the historical wildebeest habitats [9,68].

Wildebeest occupied parts of the Rift Valley extending from the Tanzania border north to Kedong Valley in Kenya and in the early 1900s, to the eastern and southern shores of Lake Naivasha [46, 69–75]. Thus, in 1902, Meinertzhagen [74] recorded wildebeest on the flats east of Lake Naivasha. Moreover, based on annual hunting returns, wildebeest were shot in Naivasha in 1906-07 [70] and in the Rift Valley in 1909-10 [71]. The Mosiro Plateau in Narok County was the probable link between the Rift and Serengeti-Mara wildebeests when the Plateau landscape was still open and had tall grass, few gullies and bushes [46]. Livestock overgrazing degraded the plateau making it impassable for wildebeest by early 1960s [46]. Part of this wildebeest population also occupied the centre and west of the Rift Valley, near Mt. Suswa, south of Naivasha in 1909 and 1910 [73]. Their distribution extended to the East Rift Wall, near Kijabe in central Kenya [76]. But, all the wildebeest populations in the northern area of the Rift Valley in Kenya became extinct before 1962 because of hunting and fencing of ranches around Lake Naivasha and in the Rift neighbouring Suswa area [46]. Wildebeest were later re-introduced on the Cresent Island in Lake Naivasha and slowly spread, or were physically moved, to other surrounding areas. Their population increased steeply in Nakuru Conservancy in the Nakuru-Naivasha region of Kenya during 1996-2015 [77]. Wildebeest were also found in several other parts of Kenya in earlier years where they have since been exterminated. In particular, returns of game animals shot on license in 1909-10, show wildebeest were found on the Mau Plateau in Narok County, the Kisii region in Southwestern Kenya, Laikipia and North Uaso Nyiro in Central Kenya, Makindu and Voi in southeastern Kenya [71].

A small wildebeest population occurs in the Ngorongoro Conservation Area in Tanzania [78,79]. It concentrates in the Crater in the dry season but disperses to areas outside the Crater in the wet season, including the Lake Natron Game Controlled Area. During the early dry season wildebeest sometimes move east from the Serengeti short grass plains into Ngorongoro Crater/Conservation Area and leave at the onset of the rains but a smaller residual population remains in the Crater during the rainy season [46,63]. Wildebeest from the NCA and the Serengeti short grass plains also migrate to the south-eastern part of the LGCA.

The Athi-Kaputiei, Amboseli, Western Kajiado and Machakos County wildebeest populations were historically part of a single large migratory population that used to range over most of the present day Kajiado County in Kenya until the 1960s before it split up into three rather distinct populations [22,26,35,76]. The western Kajiado population is currently non-migratory. The Athi-Kaputiei wildebeest population uses the Nairobi National Park during the dry season due to its reliable water supply and abundant grass and move to calve on the pastoral lands to the southeast of the park during the wet season [4,5,35,80–83]. The Athi-Kaputiei wildebeest population was centred on the Athi-Kaputiei Plains in the wet season prior to the 1920s. Historically, some Athi-Kaputiei wildebeest may have migrated south to the Amboseli ecosystem [46,67,76]. They intermingled with the then larger Amboseli population centred on the Amboseli Plains north of Kilimanjaro in the wet seasons. Both populations were migratory and moved to water in the hills and woodlands in the dry seasons and returned to the plains in the wet seasons. In very dry periods many wildebeest from both populations moved northeast and south, including into Tanzania [46].

The Athi-Kaputiei wildebeest moved north, east, and south in the wet season but spent the dry season on the Athi-Kaputiei Plains until at least 1927 [46,67,76]. The Athi-Kaputiei wildebeest population migrated as far north as the Thika River in the dry season but only few went beyond this point [46,67,76]. A resident wildebeest population north of the Thika River and another near Juja, both northeast of Nairobi in Kenya [46,75,76] went extinct. The Athi-Kaputiei wildebeest population also migrated as far north as Muranga (Fort Hall) and the Yatta Plateau, south of the Tana River [63]. De Beaton [84] recorded animal movements from the Nairobi National Park westwards towards the Ngong Hills and to the south. The Nairobi-Mombasa Road and later the park fence bordering this road interrupted the northward migration of wildlife to Nairobi, Ruiru-Thika and Ol Donyo Sapuk in the dry season. When a fence was first erected around Nairobi in the early 1900s, it killed many animals, including wildebeest [85]. A fence constructed in 1967 along the eastern side of the Ngong Hills and South of the Kiserian River and that joined the south-western corner of the Nairobi Park fence, further disrupted the dry-season migration of wildlife to the Ngong Hills [81,86].

Large areas of the Athi-Kaputiei Plains were ploughed and planted with wheat to contribute to war time food production during World War II. The wheat attracted large wildlife herds, including wildebeest, which were shot as part of crop protection [87]. This population dispersed periodically to the adjoining Machakos County, especially during droughts, but more recently, due to displacement by extreme land use changes and developments in the Athi-Kaputiei Ecosystem [5,33,35,83,88].

The migration and ranges of the Amboseli and West Kajiado wildebeest populations are described by several authors [22,25,46,63,67,75,76]. Occasional old bulls moved from the Rift near Lake Magadi area in western Kajiado to Nairobi area in the 1920s [76]. Wildlife, especially wildebeest and zebra, were also harvested in large numbers in Kajiado County (Amboseli Ecosystem) during World War II to provide meat for labour and prisoners of war; free meat for the Kamba people because of famine caused by severe drought; and after the end of the war to reduce competition with Masai livestock for forage [64,67].

In the Tarangire-Manyara Ecosystem, the migratory wildebeest occupy the Tarangire National Park in the dry season but disperse to their wet season ranges and calving grounds on the Simanjiro Plains, the Mkungunero Game Reserve, Lolkisale Game Controlled Area, Manyara Ranch, Lake Manyara National Park and adjacent game controlled areas (used mainly as hunting areas) in the wet season. OIKOS and TANAPA [37] confirmed that nine main migratory routes that Lamprey [38] had identified earlier in the Tarangire-Manyara ecosystem, were still being used during 1995-2002.

Wildebeest migrated from within protected areas in the dry season to dispersal areas outside conservation areas in the wet season in all the ecosystems but the Serengeti-Mara ecosystem where the migration occurred mostly within the protected areas (Fig. 2). Wildebeest migration has discontinued altogether in parts of the ecosystems, and reduced along a number of historical routes (Fig. 3).

**Fig 3.**
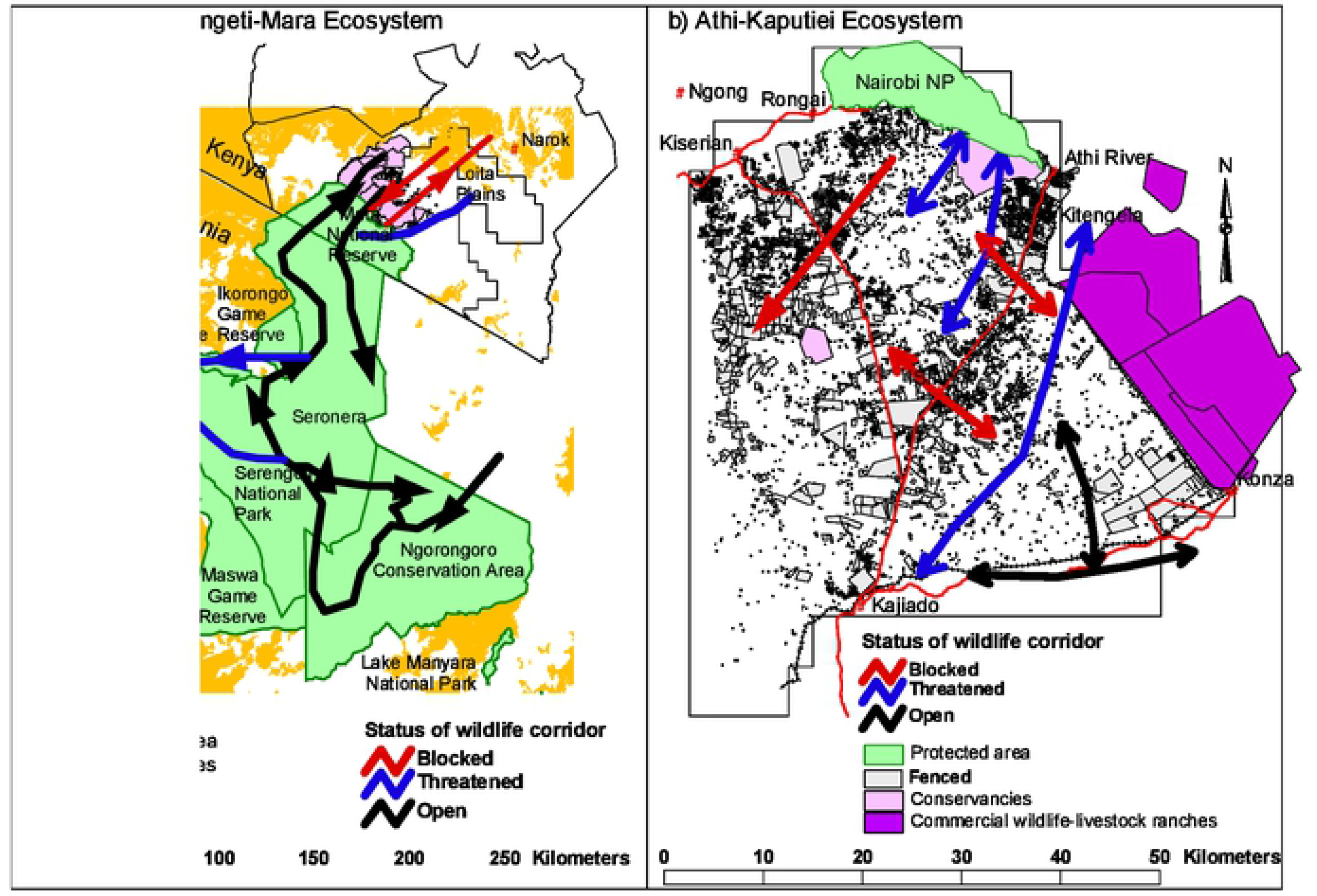
Map showing the general area occupied by the (a) Greater Serengeti-Mara, b) Athi-Kaputiei, (c) Greater Amboseli and (d) Tarangire-Manyara ecosystems. For each ecosystem, the status of routes of migratory wildebeest post-2000 in relation to the distribution of agriculture and fences is highlighted. The current wildlife conservancies (and wildlife-livestock ranches) are provided for the Masai Mara, Athi-Kaputiei and Greater Amboseli ecosystems of Kenya. Also shown are extreme land fragmentation through fences in Athi-Kaputiei and recent emergence of fences along the eastern and south eastern borders of the Mara Conservancies.

Notably, the decline and discontinuation of migration happened in four out of the five ecosystems where wildebeest migrated outside protected areas. No discontinuation of migration is reported from the Serengeti-Mara ecosystem where wildebeest migrates almost entirely within protected areas. Discontinued or currently less intensively used migration routes overlapped with agricultural and settlement expansion in the Mara and Tarangire-Manyara ecosystems and fences, settlements and roads in the Athi-Kaputiei Plains (Fig. 3). Fig. 3 does not include settlements, which is another main cause of change to the migratory routes in the study ecosystems.

Movement data (*n* = 279,718 fixes) from the GPS collared wildebeest showed the migration routes during 2010-2013 (Fig. 4). Several features of the wildebeest movements and space use are noteworthy. Wildebeest primarily used habitats outside of the protected areas in the Mara, Athi-Kaputiei and Amboseli ecosystems (> 87% of the 279,718 fixes). This emphasizes the importance of pastoral lands and community-based conservation to the protection of the three wildebeest populations. In particular, the Loita Plains wildebeest heavily used the wildlife conservancies adjoining the Masai Mara National Reserve to the north. Hence, when both the reserve and conservancies are considered, 73.4% (85,194 of 116,061 fixes) of the Loita Plains wildebeest locations fell within the conservation area boundaries. Further, one wildebeest collared in Loita Plains moved south through the LGCA to the NCA in Tanzania, covering a total of 205.4 km from its initial collaring location (Fig. 4b). This route approximates the historical migration route of the Loita wildebeest up to the 1950s [63]. This reinforces the critical importance of LGCA to Serengeti-Mara, Loita and Ngorongoro wildebeest migrations and to the ecological integrity of the Greater Serengeti-Mara ecosystem.

**Fig 4.**
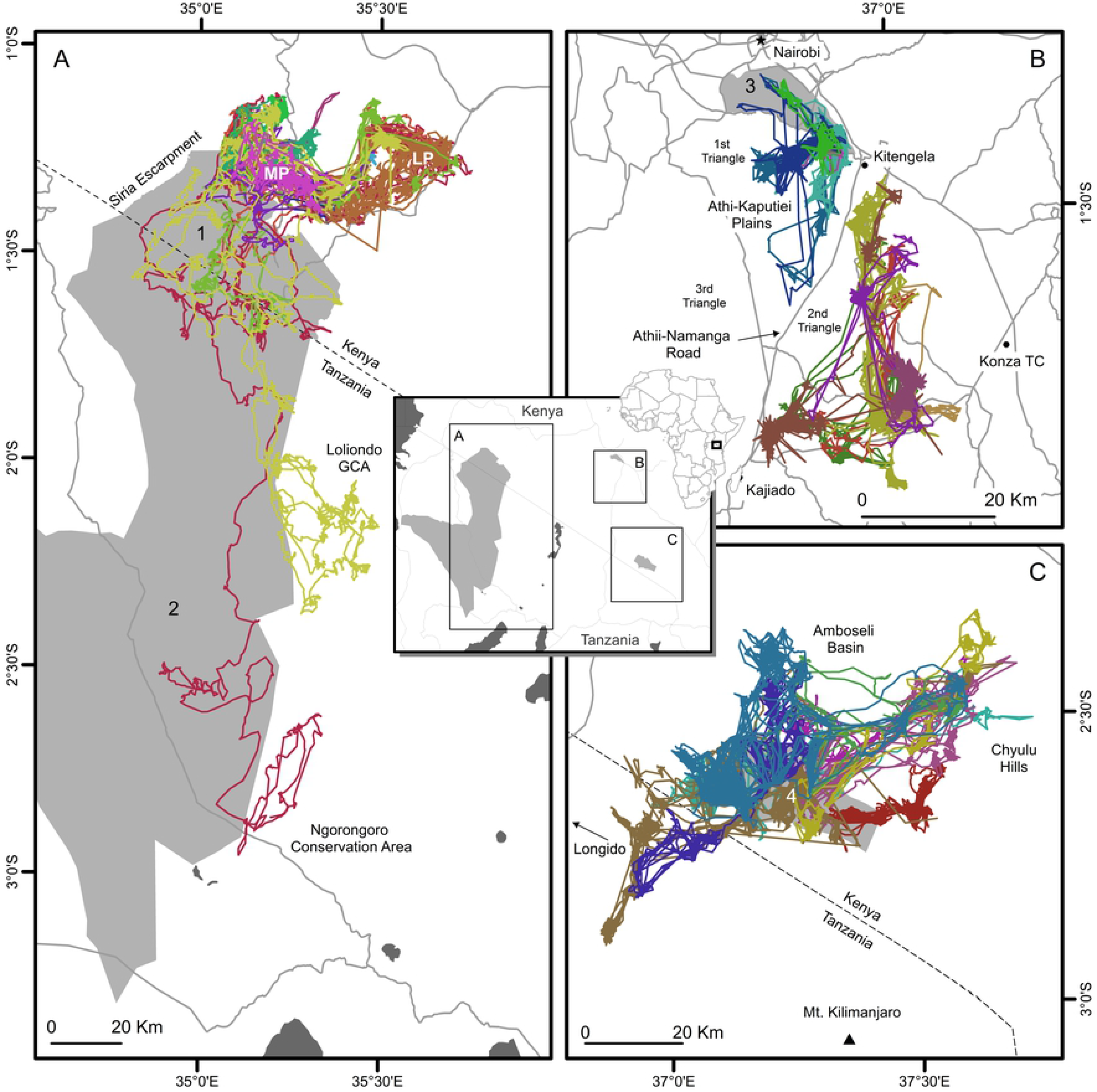
Movement tracks of GPS collared wildebeest during 2010-2013 (colored lines) in Kenya (A = Loita Plains in Masai Mara Ecosystem, B = Athi-Kaputiei Ecosystem, C = Greater Amboseli Ecosystem). Protected areas (1 = Masai Mara National Reserve, 2 = Serengeti National Park in Tanzania, 3 = Nairobi National Park, 4 Amboseli National Park) are partially obscured.

The Nairobi-Namanga tarmac road, bisecting the wet season range of the Athi-Kaputiei wildebeest, has split the population into two distinct sub-populations, concentrated on the eastern and western sides of the road (Fig. 4b). Collaring locations and direct field observations showed that no collared wildebeest crossed the tarmac road during the 2010-2013 study period. Lastly, the Amboseli wildebeest population also moved widely, including into the adjoining Longido District in Tanzania, reflecting the historical migration routes for this population [46,76]. One wildebeest collared in the Amboseli Basin travelled 6,197.8 km over 728 days during the study period. Further details on the collared wildebeest movements can be found in Stabach [88].

### Wildebeest population trends

The Serengeti-Mara wildebeest population grew steadily from 190,000 in 1957 following the veterinary eradication of rinderpest in cattle in 1962 [90], until 1977 when it stabilized with one noticeable decline during 1993, when a severe drought reduced the population from around 1.2 million to less than 900,000 animals [91] (Fig. 5a). The population has since then recovered and stabilized at around 1.3 million animals [14] though the more recent population size estimates suggest some slight upward trend (Fig. 5a). The estimated population size and standard errors and other details of the aerial surveys for all the eight study ecosystems are provided in S1-S5 Datas.

**Fig 5.**
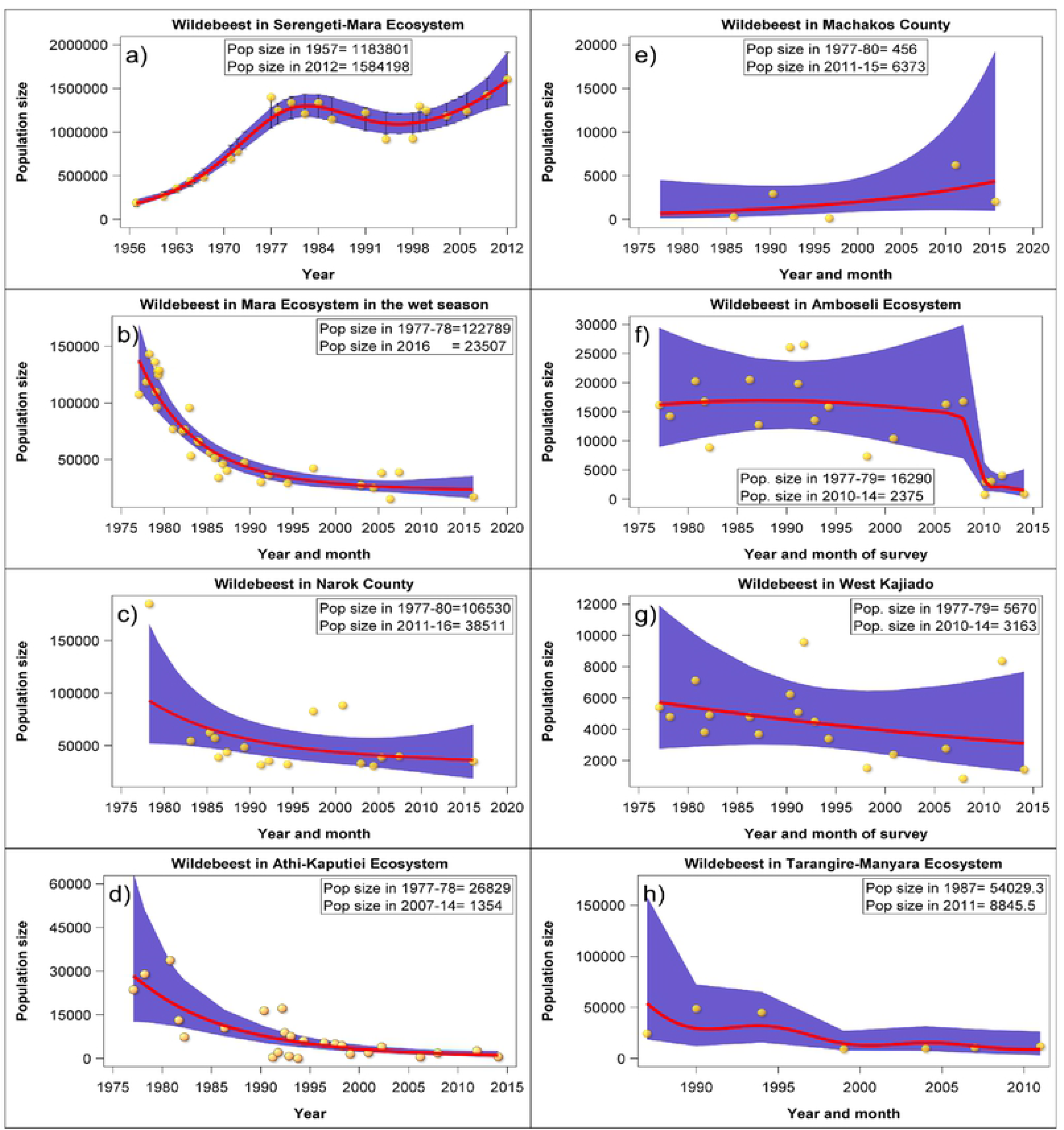
Trends in population size of migratory wildebeest populations in a) Serengeti-Mara ecosystem, b) Masai Mara ecosystem, c) Narok County in which Masai Mara is located, d) Athi-Kaputiei ecosystem, e) Machakos County, f) Greater Amboseli ecosystem, g) West Kajiado and h) Tarangire-Manyara ecosystem.

The Loita Plains wildebeest population declined steadily from about 123,930 animals in 1977-1978 to around 19,650 animals by January 2016 (Fig. 4b), a decrease of 80.9%. This decline was highly significant (Table 1). The population of the Serengeti migrants coming to the Mara ecosystem in the dry season (July-October) similarly decreased by 73.4% from 587,500 in July-August 1979 to 157,124 animals in November 2016. The dramatic decline was also evident for the Narok County wildebeest population (Table 1, Fig. 5c, S2 Data).

**Table 1.**
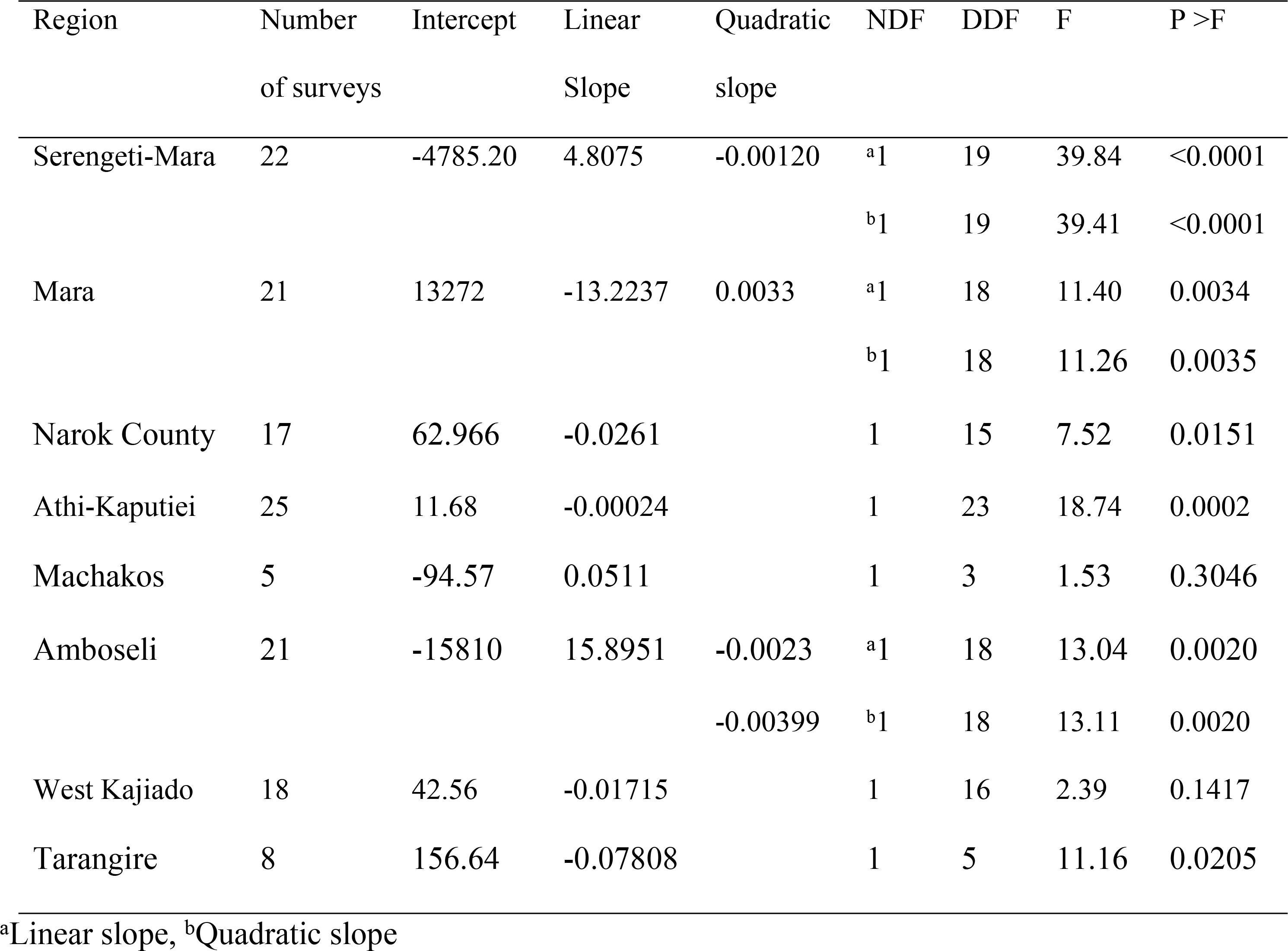
Results of the regression of wildebeest population size on year of survey. NDF and DDF are the numerator and denominator degrees of freedom, respectively.

| Region | Number<br>of surveys | Intercept | Linear<br>Slope | Quadratic<br>slope | NDF | DDF | F | P > F |
| --- | --- | --- | --- | --- | --- | --- | --- | --- |
| Serengeti-Mara | 22 | -4785.20 | 4.8075 | -0.00120 | <sup>a</sup>1 | 19 | 39.84 | <0.0001 |
|  |  |  |  |  | <sup>b</sup>1 | 19 | 39.41 | <0.0001 |
| Mara | 21 | 13272 | -13.2237 | 0.0033 | <sup>a</sup>1 | 18 | 11.40 | 0.0034 |
|  |  |  |  |  | <sup>b</sup>1 | 18 | 11.26 | 0.0035 |
| Narok County | 17 | 62.966 | -0.0261 |  | 1 | 15 | 7.52 | 0.0151 |
| Athi-Kaputiei | 25 | 11.68 | -0.00024 |  | 1 | 23 | 18.74 | 0.0002 |
| Machakos | 5 | -94.57 | 0.0511 |  | 1 | 3 | 1.53 | 0.3046 |
| Amboseli | 21 | -15810 | 15.8951 | -0.0023 | <sup>a</sup>1 | 18 | 13.04 | 0.0020 |
|  |  |  |  | -0.00399 | <sup>b</sup>1 | 18 | 13.11 | 0.0020 |
| West Kajiado | 18 | 42.56 | -0.01715 |  | 1 | 16 | 2.39 | 0.1417 |
| Tarangire | 8 | 156.64 | -0.07808 |  | 1 | 5 | 11.16 | 0.0205 |

<sup>a</sup>Linear slope, <sup>b</sup>Quadratic slope

The Athi-Kaputiei wildebeest population suffered a 95% decline in numbers from over 26,800 animals in 1977-1978 to less than 10,000 by the mid-1990s and under 3,000 animals in 2007-2014. The decline of this population has been much more dramatic in recent decades, leading to a virtual collapse of the migration (Fig. 5d). The catastrophic decline is highly significant (Table 1, S3 Data). A recent 1298% increase in Machakos County population, coincident with the decrease in the Athi-Kaputiei population (Fig. 5e), is not statistically significant likely because of a large variance in the population estimates (Table 1, S4 Data). This strongly suggests that some wildebeest migrated from the Athi-Kaputiei rather than died.

The migratory wildebeest population in the Amboseli ecosystem also declined by 84.5% from about 16,290 animals in 1977-1979 to 2,375 by 2010-2014 (Fig. 5f). The population fluctuated between 16,290 and 20,000 individuals and increased to 33,000-37,000 individuals during 1978-1986 and fell to 16,779 animals by 2007. The population declined to under 5,000 animals in 2010 following a severe drought in 2008-2009 (Fig. 5f) and has not recovered ever since. This decline is highly statistically significant (Table 1). The non-migratory wildebeest population in West Kajiado decreased by 44% from 5,700 animals in 1977-1979 to 3200 animals in 2010-2014 but this decrease is not statistically significant likely due to large variances in population size estimates (Table 1, Fig. 5g, S5 Data).

The Tarangire-Manyara population first increased from an estimated 24,399 animals in 1987 to 48,783 animals in 1990. Thereafter the population fell precipitously to 13,603 animals by 2016 without signs of recovery (Fig. 5h). This extreme population decline is statistically significant (Table 1) despite the large variances in the population estimates (S5 Data).

## Discussion

### Wildebeest movements and migratory routes

Animal movement depends on individual fitness and is essential for accessing favoured resources, finding potential mates and escaping deteriorating habitat conditions [92]. As expected, the GPS collared wildebeest moved more, in virtually all measured aspects, in Amboseli, the least productive and least anthropogenically disturbed of the three Kenyan ecosystems, than in the Loita Plains and Athi-Kaputiei. The productivity of Amboseli grasslands has reduced even further in recent years [93,94], apparently forcing wildebeest to move over larger areas in search of food in the dry season [95]. Wildebeest surprisingly moved less in Athi-Kaputiei than in either the Amboseli or Loita Plains even though the Loita Plains had the greatest availability of resources of the three landscapes. This is unexpected even if the wildebeest decline in the Athi-Kaputiei has reduced intraspecific competition and the need to move to locate resources. High livestock density likely heightens interspecific competition with wildebeest for resources and thus could force wildebeest to move more in Athi-Kaputiei. The reduced wildebeest movements in the Athi-Kaputiei landscape therefore reflect its high degree of anthropogenic disturbance and truncation [5], preventing needed further movement [88]. It follows that resource availability and anthropogenic disturbance determine wildebeest movements. Consequently, because wildebeest occur primarily outside protected areas, except in the Serengeti-Mara, controlling the rate and type of anthropogenic change in these areas is crucial to maintaining the long-term viability of their populations and migrations.

### Wildebeest population declines

Migratory wildebeest population size and their routes declined in all the five ecosystems except the Serengeti-Mara. The declines are related to expansion of agriculture, settlements, fences and roads that progressively occlude wildebeest grazing resources and migratory routes (Table 2). Even though it was not possible to formally test if these processes caused the declines, literature review, interviews and collared wildebeest movements, suggest that they are all important. In all the four ecosystems where they are declining, agricultural encroachment excludes wildebeest from part of their seasonal ranges. Notably, irrigated agriculture encroached the swamps that ring the base of Mt. Kilimanjaro, denying wildebeest access to their critical dry season dispersal areas in Amboseli [24,25,96]. Settlements also interfere with wildebeest movements in the Mara [31,34,52], Tarangire-Manyara [15,18] and Athi-Kaputiei [4,5,35] by blocking their migratory routes and access to resources. Further, although wildebeest avoid anthropogenic disturbances [97], they are attracted to short grass created by livestock grazing outside protected areas on pastoral lands with moderate densities of pastoral settlement and livestock [21,98].

**Table 2.** Summary of the processes likely associated with the declining migratory wildebeest populations and patterns in the East African rangelands.

| Processes | *Serengeti-Mara | Masai<br>Mara | Athi-Kaputiei | Greater<br>Amboseli | Tarangire-Manyara |
| --- | --- | --- | --- | --- | --- |

| **Direct interferences/causes** |  |  |  |  |  |
| --- | --- | --- | --- | --- | --- |
| Agricultural encroachment | - | +++ | + | +++ | +++ |
| Fencing | - | ++ | +++ | ++ | - |
| Settlements | + | ++ | +++ | ++ | ++ |
| Urbanization | - | + | +++ | - | - |
| Roads & Infrastructural developments | + | ++ | +++ | + | ++ |
| Poaching by increasing human populations | ++ | + | ++ | + | ++ |
| Competition with livestock for forage,<br>water and space | + | +++ | +++ | +++ | +++ |
| Declining drinking water supply and<br>quality | + | ++ | +++ | ++ | + |
| **Drivers** |  |  |  |  |  |
| Human population increase | + | ++ | +++ | ++ | +++ |
| Land tenure change | + | ++ | +++ | +++ | + |
| Land subdivision | - | +++ | +++ | ++ | ++ |
| Settlement policies | ++ |  | +++ |  | +++ |
| Wildlife conservation and management<br>policies | +++ | +++ | +++ | +++ | +++ |
| Wildlife management institutions | - | +++ | +++ | ++ | ++ |
| Wildlife markets or benefits to landowners | - | +++ | +++ | +++ | + |

‡+++ High importance; ++ Important; + less importance; - not important. Source: Interviews with resident researchers [4,5,15,18,111].

Land fragmentation through fencing, roads and settlements primarily exclude wildebeest from their grazing ranges in the Athi-Kaputiei ecosystem [5] (Table 2). In Kitengela, a major part of the Athi-Kaputiei Plains adjoining Nairobi National Park, fenced land parcels have spread throughout the range of wildlife and movements of people, livestock, dogs and vehicles harass wildlife [4,5,35]. Fences impede wildebeest movements between the Nairobi Park and the Athi-Kaputiei Plains [4,5,35]. Similarly, the Nairobi-Namanga road has effectively truncated the ecosystem, splitting the Athi-Kaputiei population into two separate sub-populations [89]. The Athi-Kaputiei landscape is also fragmented and degraded by large, un-rehabilitated mines, mining waste, unregulated development, commercial charcoal burning and sand harvesting, all of which restrict wildebeest habitats and obstruct their migratory routes. Invasive weeds are also spreading in the rangelands and at abandoned settlement sites in Athi-Kaputiei, degrading wildebeest habitats [5,35,51,99]. Fences [100,101] are also increasing rapidly in the Mara, including in the Loita Plains, following land subdivision and privatization of land ownership. The loss of connectivity restricts the mobility and flexibility of migratory wildebeest, especially during droughts when heavy mortality can result where wildebeest access to water and food is blocked [102–104]. The risk of outbreaks of zoonotic diseases and population declines can also increase if ungulate migrations are curtailed by degraded habitats yet climate change increases the frequency and severity of droughts [105]. Climate change may amplify the frequency of outbreaks of zoonotic diseases by modifying host and vector population characteristics that control pathogen transmission, including concentration in key resource areas, population density, prevalence of infection by zoonotic pathogens, and the pathogen load in individual hosts and vectors [106,107]. Also, calving wildebeest transmit bovine malignant catarrhal fever (BMCF) virus to livestock where the two species co-occur, causing livestock losses [20]. The risk of transmitting the BMCF virus is elevated where habitat loss and degradation force livestock and wildebeest to use the same areas.

Another leading cause of wildebeest decline is poaching, which removes 6-10% of the Serengeti-Mara wildebeest annually [108,109]. Poaching is also common in the other ecosystems, including the Mara [34] and Athi-Kaputiei [35]. The status and threats facing the five ecosystems with migratory wildebeest populations in East Africa are summarized in Table S2.

Additional factors that adversely affect access of migratory wildebeest to critical habitats, food and water include human population expansion, land subdivision and privatization of land tenure, development of urban centres and intensification of land use following sedentarization of formerly semi-nomadic pastoralists [4,26,31,33–35]. Why is the Serengeti-Mara wildebeest population stable while the other populations are declining? Moreover, given that the Serengeti-Mara ecosystem protects nearly 1.5 million wildebeest, why should we worry about conserving the other smaller wildebeest populations? First, the Serengeti-Mara populations are not declining because over 80% of the wildebeest live in the large and relatively well-protected ecosystem. Second, it is important to conserve the smaller populations in other areas for at least three reasons. a) Some of the other areas support populations of wildebeest belonging to a different subspecies from that found in Serengeti-Mara. b) Migratory wildebeest provide important ecosystem services, even at low densities, such as promoting calf survival among other ungulate species by reducing predation pressure when present in an area [112]. c) Wildebeest migrations are a magnificent spectacle and thus can provide significant tourism revenue opportunities in specific areas.

### Land-use change and poaching as causes of wildebeest population declines

Land use change, particularly expansion of agriculture, settlements and fences and commercial charcoal production linked to human population growth degrade and reduce wildlife habitats [5,26]. In the Athi-Kaputiei ecosystem, expansion of the neighbouring Nairobi Metropolis, urbanization in the ecosystem plus relatively lower land prices compared with Nairobi, strongly drive land use change [114]. Development of new industries, businesses and infrastructure attract more people from Nairobi and elsewhere to the Athi-Kaputiei [4,83]. In Amboseli and Western Kajiado, commercial charcoal production is causing widespread deforestation of wildlife habitats [115].

Agriculture, particularly large-scale commercial cultivation, is a leading cause of habitat loss for the migratory wildebeest. Previously, mainly outsiders practiced cropped agriculture in the study ecosystems, but the Masai have recently started cultivating next to their settlements [116,117]. Widespread adoption of subsistence agriculture in small plots right around a household’s compound can threaten wildebeest populations migrating outside protected areas [118]. Remarkably, nearly 500 km^2^ of natural vegetation in the Loita Plains were converted to wheat farms and other uses between 1975 and 1995 [67] and even more has been converted in recent years [119,120]. In Tanzania, people moved into, and cultivated for several years, parts of Game Controlled Areas or Open Areas, such as the LGCA, which had functioned much like game reserves in the past, interfering with wildlife migrations.

In the Tarangire-Manyara ecosystem, about 710 km^2^ of land was converted from rangelands to farms between 1984 and 2000 [15], cutting-off large portions of forage and dispersal areas and blocking routes traditionally used by migratory wildebeest. Villagization promoted by government settlement policies in Tanzania is another key driver of land conversion to agriculture in the Tarangire-Manyara ecosystem [15] and western Serengeti [14].

Wildebeest prefer land suitable for agriculture [121] and therefore face a high risk of displacement by agriculture and competition with livestock for space, forage and water in pastoral lands. Such high potential lands tend to generate higher economic returns from cropping than from livestock or conservation [114,119]. Land users are thus likely to opt for cultivation rather than conservation thereby accentuating encroachment of agriculture and wildebeest population declines. But once land is cultivated, it is difficult to restore to its former rangeland status where returns from agriculture overwhelm those from either livestock or wildlife [119]. Even where wildlife tourism benefits are competitive with those from agriculture, the benefits often accrue to the rich so that the poor land owners, who bear the burden of supporting wildlife on their lands, typically receive meagre benefits [122]. This calls for schemes for more equitable sharing of wildlife benefits [116].

Land tenure change from group ranches to private ownership is another important driver of land use change in Masailand in Kenya [116]. The land sub-divisions and individualization of tenure associated with fencing in Masai Mara, Athi-Kaputiei and Amboseli ecosystems amplify habitat fragmentation and interfere with the migratory wildebeest [4,5,123].

Poaching is associated with increasing human population size and resource use intensity [108–110]. On commercial wheat farms in the Mara, poaching is very common (R. Lamprey, pers comm), especially far from pastoral settlements, because pastoralists often discourage poaching. Poaching is also common inside the protected areas in the Mara and Serengeti [34,109,110]. In the Athi-Kaputiei, poachers killed many wildebeest by running them up against fences [35].

### Ways to make land use compatible with wildlife conservation

Human population explosion, unplanned urbanization, settlements, cultivation and other developments pose unprecedented challenges to conservation and maintenance of migrations as the spaces available for wildlife and their habitats shrink, leading to population declines. It is thus important to conserve spatially extensive migratory systems while balancing human and wildlife needs. In Kenya, wildlife conservancies are expanding conservation areas for wildlife beyond the state-owned parks and reserves onto land owned privately by local communities or individuals who benefit by receiving land rents and job opportunities [34,113].

It is primarily tourism income that pays for conservancy land leases and management in Kenya. Thus, the success of the common conservancy model in Kenya is contingent upon sustainable wildlife tourism making it worthwhile for landowners to allow conservancies to be set up on their lands. This conservancy model can thus only be viable in areas with low tourism potential if tourism revenue is supplemented with other revenue streams.

Nevertheless, wildebeest can and do benefit from community-based wildlife conservation endeavours where wildlife conservancies have been established on private and communal rangelands, including in areas of high rainfall [14,25,124,125]. By 2015, 178 wildlife or mixed livestock-wildlife conservancies had been established across Kenya [51] and new ones continue to be established on private and communal lands in Masai Mara, Amboseli, Athi-Kaputiei and Machakos (Tables S3 and S4), Naivasha-Nakuru and other parts of Kenya [26,34,77,125,126]. The total area of wildlife conservancies and ranches in Kenya’s rangeland counties by 2017 was 54,265 km^2^ of which Narok, Kajiado and Machakos counties that support wildebeest populations had set aside 2,219, 2,837 and 463 km^2^, respectively (KWCA, Unpublished data, https://kwcakenya.com/).

What makes the conservancies so popular with local communities is that they also protect land rights; create jobs; provide income to communities through tourism; and provide increased security for people, livestock and wildlife [51]. Conservancies are crucial for wildlife conservation because all state protected areas cover only about 10% of Kenya’s land surface (an additional 10.7% is in conservancies, benefiting close to 700,000 people nationally) and 70% of these areas are found in the rangelands of Kenya. Moreover, about 65% of Kenya’s wildlife are found outside the protected areas [127]. As limited public land constrains expansion of public protected areas, the private and communal conservancies are crucial for expanding the space for wildlife in Kenya. The conservancies are promoting positive attitudes towards wildlife and restoration of degraded rangelands by regulating livestock grazing, restricting settlements and other developments. They act as buffers for parks and reserves, besides offering increased protection to wildlife, enabling many wildlife species to increase within conservancies [77,126,128].

Effective wildlife conservation would require permanent conservancies, land purchases or conservation easements on land used by wildlife. In the Kenya wildlife conservancies, landowners typically amalgamate adjacent individual plots to create large, viable game viewing areas. They then broker land lease agreements with a coalition of commercial tourism operators under institutional arrangements modelled in the form of payments for ecosystem services [125]. There is a strong interest in this wildlife conservancy model in Kenya. Thus, starting with only two conservancies in 2005-2006 covering 145.76 km^2^, there were eight conservancies covering about 1000 km^2^ by 2010 [125] and 10 conservancies by 2016 (Table S3). The Mara conservancies covered 1355 km^2^ by 2018 and are expanding rapidly. The development of the Mara conservancies has helped partially unblock the movements of migratory wildebeest between the Mara Reserve and the Loita Plains.

In certain areas, such as the Athi-Kaputiei, land owners are paid conservation land lease fees since 2000 to keep land open for use by wildlife and livestock, not building fences and for collecting poachers’ snares [35,129–131]. The cost of financing such land leases over large areas year after year would, however, require creating conservancies able to maintain viable conservation enterprises, such as a vibrant tourism industry, to ensure their long-term sustainability. The benefits derived from such enterprises would be an important incentive for the landowners to continue keeping their land open for use by wildlife and desisting from other uses incongruent with conservation. The changes taking place in Athi-Kaputiei are, however, so dramatic and fast that unless these conservation efforts are undertaken immediately, the opportunity to save even the very few remaining and most critical portions of this once magnificent ecosystem, is highly likely to be lost for good.

In Tanzania, various conservation initiatives have been launched to protect the remaining migratory routes and dispersal ranges beyond the borders of protected areas. These include reducing illegal hunting and livestock grazing in Manyara Ranch, recently converted to a private Conservancy. Provision of artificial water holes in the Manyara Ranch Conservancy keeps migratory wildebeest and zebra in the vicinity of the Conservancy until late in the dry season. On the communal grazing lands, initiatives have been launched to enhance the wildlife benefits going to the local communities. In Simanjiro Plains, hunting companies, tour operators and conservation organizations have teamed up together to pay for conservation land lease fees to community members to refrain from farming or expanding settlements into critical areas of communal grazing lands [132]. Certificate of Customary Right of Occupancy (CCRO) agreements are also being used to protect grazing ranges for wildlife and pastoral livestock, including in areas neighbouring migratory corridors. One such CCRO was established in Selela Village situated north of Manyara and includes an important but narrow corridor [133]. Other initiatives include establishment of Wildlife Management Areas (WMAs) two of which were recently established to the north and west of Tarangire National Park [18]. The WMAs not only protect communal land but also reduce the incentive for poaching by distributing tourism revenue to the local communities. The Tarangire National Park and the Tanzania Wildlife Authority are also supporting community game rangers to intensify anti-poaching patrols in the WMAs and Manyara Ranch Conservancy and among villages in the Simanjiro Plains.

### Wildlife conservation initiatives and gaps in wildlife policies, institutions and markets in Kenya and Tanzania

What else can be done to stop the declines and allow migratory wildebeest access to at least the few remaining critical portions of their former habitats? A significant challenge to wildlife conservation in East Africa remains incoherent government development policies that promote incompatible land uses, such as promoting cultivation in pastoral rangelands occupied by wildlife to combat food insecurity while also promoting wildlife-based tourism in the same areas. Such policies should be harmonised to minimize the adverse impacts on wildlife conservation of incongruent land uses in pastoral rangelands. Another weakness of the wildlife policy in Kenya is that the state owns all wildlife whereas land owners in the rangelands do not have access to or user rights over wildlife. The land owners do not get any compensation for the opportunity cost of supporting wildlife on their private lands nor for wildlife damage to their private property, thus fuelling indifference or hostility towards wildlife. There is also no public institution specifically charged with conserving and managing wildlife on the private lands. Although these shortcomings are well documented [114,134] and have partly been addressed by the Wildlife Conservation and Management Act 2013 [135] and the National Wildlife Strategy [136], the Act should be fully implemented to address these glaring policy, institutional and market deficiencies.

In Tanzania, several national initiatives are being undertaken to restructure the institutions that manage the wildlife sector in order to contain a spiralling poaching crisis. Key among these is the dissolution of the former Wildlife Division (WD) that used to manage all the Game Reserves and Game Controlled Areas, including overseeing all wildlife in village lands (i.e., WMAs), and its reconstitution as the Tanzania Wildlife Authority (TAWA) in October 2015. TAWA is empowered and better funded compared with its predecessor, the WD, to improve the management of the wildlife areas under its jurisdiction.

The second is the re-organization of the entire wildlife sector in the country into para-military style organizations to intensify the fight against run-away poaching in protected and unprotected areas, most especially in game reserves. Because many game reserves and game controlled areas share open borders with national parks, wildlife population declines due to poaching are occurring even inside the national parks. But, to be successful in curbing poaching, these efforts should be accompanied with enhanced economic incentives to communities neighbouring wildlife areas or sharing land with wildlife to discourage poaching and destruction of wildlife habitats. Tanzania is also working on expanding the Serengeti National Park by adding to it about 1500 km^2^ from the Loliondo Game Controlled Area to the east and extending the western side of the park to reach the shores of Lake Victoria.

A major initiative in both Tanzania and Kenya is the development of national policies on wildlife corridors, dispersal areas, buffer zones and migratory routes to promote habitat connectivity [137,138]. Regional initiatives linking the two countries are, however, needed to foster close cooperation between Kenya and Tanzania in conserving the trans-boundary wildebeest migrations and implementing regional and international conservation conventions and treaties. Such initiatives should include harmonization of policies, legal and regulatory frameworks for the conservation of wildlife and other species involved in trans-boundary migrations.

## Conclusions

Migratory wildebeest populations in four out of five key ecosystems in East Africa are under severe threats and two populations are on their way to total collapse if the trends are left to continue unabated. Such collapse in migratory wildlife population in East Africa has been documented for zebra and Thomson’s gazelle populations that used to migrate between Lakes Nakuru and Elementaita and Baringo regions of Kenya [76,77,139] that went extinct because of fences and uncontrolled shooting [85]. The migration of the Athi-Kaputiei wildebeest to Nairobi National Park had also virtually collapsed by 2011 [35]. Recent surveys in the park show that the wildebeest involved in this migration remained under 350 animals from 2012 to 2015 [5]. Agricultural encroachment, settlements, poaching, roads and fencing are the major proximate threats responsible for the extreme wildebeest losses and degradation of their habitats as they directly kill, displace, or reduce wildebeest access to forage, water and calving areas. The fundamental causes of wildebeest population declines seem to be expanding unplanned land use developments driven by human population growth; poaching, policy, institutional and market deficiencies. Consequently, the Kenyan and Tanzanian governments need to strongly promote and lead the conservation of the remaining key wildebeest habitats, migration corridors and populations to ensure their continued access to grazing resources in these rangelands. More wildlife conservancies or management areas should be established to protect migratory routes or corridors, buffer zones, dispersal areas and calving grounds for the species. Land use and development planning should be enhanced and gaps in wildlife policies, institutions and markets addressed. Where migration occurs across international boundaries, such as in the Serengeti-Mara, Loita Plains and Amboseli ecosystems, wildlife policies, land use plans, conservation and management goals should be harmonized to ensure the long-term survival of migratory species and the sustainability of the rangelands upon which they depend. All areas currently under protection should ideally have binding legal restrictions on future developments to minimize their vulnerability to future changes. The various conservation initiatives should be coordinated spatially and across bureaucratic lines to enhance their effectiveness.

## Acknowledgements

We thank the Tanzania Wildlife Research Institute (TAWIRI) and the Directorate of Resource Surveys and Remote Sensing of Kenya (DRSRS) for permission to use their aerial survey data. JOO was supported by a grant from the German National Research Foundation (DFG; Grant # OG 83/1-1). MYS was supported by the Pathways to Resilience in Semi-Arid Economies (PRISE, Project # 107643-001). This project has received funding from the European Union’s Horizon 2020 research and innovation programme under grant agreement No 641918 through the AfricanBioservices Project. National Science Foundation (NSF) DEB Grant #0919383) supported this work through the project: Wildebeest Forage Acquisition in Fragmented Landscapes under Variable Climates.

## Supporting information

**Table S1. The field data form used to collect information for ground truthing historical information on land use and cover changes in the Tarangire-Manyara ecosystem of Tanzania.** We uploaded the coordinates and their Ids into a Global Positioning System (GPS) and used them to locate sampling points. We collected information on the following attributes for each sampling point. (1) Was there agriculture at the location in 1984, 2000 or 2006-2007? (2) If yes, was agriculture practiced on small or large scale? (3) Was agriculture irrigated or not? (4) When did agriculture start? (5) Photographs of the sampling point. (6) Crop types cultivated at each sampling location. (7) We empirically assessed the change type we had identified during initial image interpretation in the office and assigned change codes A, B, C or D described in the table to the observed changes. (8) General comments.

## Supporting information

**Table S1**.

**Table S2. The five ecosystems with migratory wildebeest populations in East Africa, their current status and earlier studies**.

**Table S3. Wildlife conservancies in Masai Mara, their names, size, number of landowners that pooled land to form the conservancy, tourist camps, tourist beds, rangers and scouts and jobs created by each conservancy and year of establishment**.

**Table S4. Wildlife conservancies or ranches, their names and sizes in Machakos Plains (adjoining the Athi-Kaputiei), Athi-Kaputiei and Greater Amboseli ecosystem.** The total area covered in each ecosystem is 347.0, 40.6 and 1046.5 km^2^ in the Machakos Plains, Athi-Kaputiei and Greater Amboseli ecosystems, respectively.

**S1 Data. The estimated population size and standard errors and other details of the aerial surveys of wildebeest for the Serengeti-Mara (1957-2012) and Tarangire-Manyara (1987-2016) ecosystems.**

**S2 Data. The estimated population size and standard errors and other details of the aerial surveys of wildebeest for the Masai Mara Ecosystem (1977-2016) ecosystems.**

**S3 Data. The estimated population size and standard errors and other details of the aerial surveys of wildebeest for the Athi-Kaputiei Ecosystem (1977-2014).**

**S4 Data. The estimated population size and standard errors and other details of the aerial surveys of wildebeest for the Machakos County (1977-2015)**.

**S5 Data. The estimated population size and standard errors and other details of the aerial surveys of wildebeest for the Greater Amboseli (1977-2014) and Western Kajiado (1977-2014) ecosystems.**

## References

1. Bolger DT, Newmark WD, Morrison TA, Doak DF (2008) The need for integrative approaches to understand and conserve migratory ungulates. Ecology Letters 11: 63–77.

2. Harris G, Thirgood S, Hopcraft JGC, Cromsigt JPG, Berger J (2009) Global decline in aggregated migrations of large terrestrial mammals. Endangered Species Research 7: 55–76.

3. Berger J (2004) The Last Mile: How to sustain long-distance migrations in mammals. Conservation Biology 18: 320–331.

4. Reid RS, Gichohi H, Said MY, Nkedianye D, Ogutu JO, Kshatriya M, Kristjanson P, Kifugo SC, Agatsiva JL, Andanje SA, Bagine R (2008) Fragmentation of a peri-urban savanna, Athi-Kaputiei plains, Kenya. In: Fragmentation in Semi-arid and Arid Landscapes;Consequences for Human and Natural Systems (eds KA Galvin, RS Reid, RH Behnke, NT Hobbs), pp. 195–224. Springer.

5. Said MY, Ogutu JO, Kifugo SC, Makui O, Reid RS, de Leeuw J (2016) Effects of extreme land fragmentation on wildlife and livestock population abundance and distribution. Journal of Nature Conservation 34: 151–164.

6. Fryxell JM, Greever J, Sinclair ARE (1988) Why are migratory ungulates so abundant? American Naturalist 131: 781–98.

7. Hopcraft JGC, Holdo RM, Mwangomo E, Mduma SAR, Thirgood SJ, Borner M, Sinclair ARE (2015) Why are wildebeest the most abundant herbivore in the Serengeti ecosystem? Serengeti IV: sustaining biodiversity in a coupled human-natural system, 125.

8. Sinclair ARE (1995) Serengeti past and present. Pp. 3–30 In: Serengeti II. Sinclair, ARE Arcese P (Eds). Chicago University Press, Chicago.

9. Ottichilo WK, de Leeuw J, Prins HHT (2001) Population trends of resident wildebeest Connochaetes taurinus hecki (Neumann) and factors influencing them in the Masai Mara ecosystem, Kenya. Biological Conservation 97: 271–282.

10. Serneels S, Lambin EF (2001). Proximate causes of land-use change in Narok District, Kenya: a spatial statistical model. Agriculture, Ecosystems Environment 85: 65–81.

11. Wolanski E, Gereta E, Borner M, Mduma SAR (1999) Water, migration and the Serengeti ecosystem. American Scientist 87: 523–526.

12. McNaughton SJ (1990) Mineral nutrition and seasonal movements of African migratory ungulates. Nature 345: 613–615.

13. Fryxell JM, Sinclair ARE. (1988) Causes and consequences of migration by large herbivores. Trends in Ecology and Evolution 9: 237–241.

14. Thirgood S, Mosser A, Tham S, Hopcraft JGC, Mwangomo E, Mlengeya T, Kilewo M, Fryxell J, Sinclair ARE, Borner M (2004) Can Parksprotect migratory ungulates? The case of the Serengeti wildebeest. Animal Conservation 7: 113–120.

15. Msoffe FU, Kifugo SC, Said MY, Neselle M, van Gardingen P, Reid RS,Ogutu JO, Herrero M and de Leeuw J (2011) Drivers and impacts of land-use change in the Masai-Steppe of Northern Tanzania; a ecology-socio-political analysis. Land Use Science 6: 261–281.

16. Homewood K, Lambin EF, Kariuki A, Kikula I, Kivelia J, Said MY, Serneels S, Thompson M (2001) Long-term changes in Serengeti-Mara wildebeest and land cover: pastoralism, population or policies? Proceedings of the NationalAcademy of Science 98: 12544–12549.

17. Morrison TA, Bolger DT (2012) Wet season range fidelity in a tropical migratory ungulate. Journal of Animal Ecology 81: 543–552.

18. Morrison TA, Link WA, Newmark WD, Foley CA, Bolger DT (2016) Tarangire revisited: Consequences of declining connectivity in a tropical ungulate population. Biological Conservation 197: 53–60.

19. Foley C, Foley L (2015) Wildlife trends and status of migratory corridors in the Tarangire Ecosystem, Ed. TP Wildlife Conservation Society, Arusha.

20. Bedelian C, Nkedianye, D, Herrero M (2007) Masai perception of the impact and incidence of malignant catarrhal fever (MCF) in southern Kenya. Preventive Veterinary Medicine 78: 296–316.

21. Reid RS (2012) Savannas of our birth: People, wildlife, and change in East Africa. Univ of California Press, Berkeley, CA.

22. Estes RD, East R (2009) Status of the wildebeest (Connochaetes taurinus) in thewild 1967-2005. Wildlife Conservation Society.

23. Pennycuick L (1975) Movements of the migratory wildebeest population in the Serengeti area between 1960 and 1973. African Journal of Ecology 13: 65–87.

24. Western D (1975) Water availability and its influence on the structure and dynamics of a savannah large mammal community. African Journal of Ecology 13: 265–286.

25. Western D (1982) Amboseli National Park: Enlisting Landowners to Conserve Migratory Wildlife. Ambio 5: 302–305.

26. Ogutu JO, Piepho H-P, Said MY, Kifugo SC (2014) Herbivore Dynamics and Range Contraction in Kajiado County Kenya: Climate and Land Use Changes, Population Pressures, Governance, Policy and Human-wildlife Conflicts. Open Ecology Journal 7: 9–31.

27. Lamprey HF (1964) Estimation of large mammal densities, biomass and energy exchange in the Tarangire Game Reserve and the Masai Steppe inTanganyika. East African Journal of Ecology 2: 1–46.

28. Borner M (1985) The Increasing Isolation of Tarangire National Park. Oryx 19: 91–96.

29. Kahurananga J, Silkilwasha, F (1997) The migration of zebra and wildebeest between Tarangire National Park and Simanjiro Plains, northern Tanzania, in 1972 and recent trends. African Journal of Ecology 35: 179–185.

30. OIKOS (2002) Analysis of Migratory movements of large mammals and their interactions with human activities in the Tarangire area Tanzania, as a contribution and sustainable development strategy: Tarangire-Manyara Conservation Project (TCMP) Final Project Report. Istituto Oikos and University of Milan, Italy in collaboration with Tanzania National Parks.

31. Ogutu JO, Owen-Smith N, Piepho H-P, Said MY (2011) Continuing wildlife population declines and range contraction in the Mara region of Kenya during 1977-2009. Journal of Zoology 285: 99–109.

32. Veldhuis MP, Ritchie ME, Ogutu JO, Beale C, Estes A, Hopcraft JGC, Morrison TA, Mwakilema W, Ojwang GO, Parr CL, Probert J, Wargute PW, Olff H (2019) The Serengeti squeeze: cross-boundary human impacts compromise an iconic protected ecosystem. Science (In Revision).

33. Ogutu JO, Piepho H-P, Said MY, Ojwang GO, Njino LW, Wargute PW, Kifugo SC (2016) Extreme wildlife declines and concurrent increase in livestock numbers in Kenya: What are the causes? PLoS ONE 10(8): e0133744. doi:10.1371/journal.pone.0133744.

34. Ogutu JO, Piepho H-P, Dublin HT, Bhola N, Reid RS (2009) Dynamics ofMara-Serengeti ungulates in relation to land use changes. Journal of Zoology 278: 1–14.

35. Ogutu JO, Owen-Smith N, Piepho H-N, Said MY, Kifugo SC, Reid RS, Gichohi H, Kahumbu P, Andanje S (2013) Changing WildlifePopulations in Nairobi National Park and Adjoining Athi-Kaputiei Plains:Collapse of the Migratory Wildebeest. Open Conservation Biology Journal 7: 11–26.

36. Msoffe FU, Said MY, Ogutu JO, Kifugo SC, de Leeuw J, van Gardingen P, Reid RS (2011) Spatial correlates of land-use changes in the Masai-Steppe of Tanzania: Implications for conservation and environmental planning. International Journal of Biodiversity Conservation 3: 280–290.

37. Tarangire Conservation Project (TCP) (1997) Analysis of migratory movements of large mammals and their interactions with human activities in the Tarangire area, Tanzania, as a contribution to a conservation and sustainable development strategy. Final report, pp.217. University of Milan and Instituto Oikos, Italy, in collaboration with Tanzania National Parks (TANAPA).

38. Lamprey HF (1963) on the ecological separation of the large mammal species in the Tarangire Game Reserve, Tanganyika. East African Wildlife Journal 1: 63–92.

39. Msoffe F U, Mturi FA, Galanti V, Tosi W, Wauters LA, Tosi G (2007) Comparing data of different survey methods for sustainable wildlife management in hunting areas: the case of Tarangire–Manyara ecosystem, northern Tanzania. European Journal of Wildlife Research 53: 112–124.

40. Norton-Griffiths M (1978) Counting Animals. A series of handbooks on techniques currently used in African Wildlife ecology. Hand book No.1; Second Edition, African wildlife Foundation, Nairobi, Kenya.

41. Grunblatt L, Said MY, Warugute R (1996) National Rangeland Report. Summary of population estimates of wildlife an livestock. DRSRS. Nairobi, Kenya, Ministry of Planning and National Development.

42. Woodworth B, Farm B (1996) Tanzania Wildlife Conservation Monitoring: Procedure Manual. Frankfurt Zoological Society, Arusha, Tanzania.

43. Stewart DRM, Zaphiro DRP (1963) Biomass and density of wild herbivores in different East African habitats. Mammalia 27: 483–496.

44. Pearsall MH (1957) Report on an Ecological Survey of the Serengeti National Park Tanganyika. Oryx 4: 71–136.

45. Darling (1960) An ecological reconnaissance of the Mara Plains in Kenya Colony. Wildlife Monographs 5, 41pp.

46. Talbot LM, Talbot MH (1963) The wildebeest in western Masailand, East Africa. Wildlife Monographs 12, 88p.

47. Western D, Nightingale DLM (2003) Environmental change on the vulnerability of pastoralists to drought: the Masai in Amboseli, Kenya. In: Africa Environmental Outlook: Human Vulnerability to Environmental Change. Earthprint on behalf of the United Nations Environmental Program. London. Available at: http://oceandocs.net/bitstream/1834/436/1/Amboseli_Masai.pdf

48. Africover land classification. Available at http://www.fao.org/3/a-bd854e.pdf

49. Okello MM, D’amour DE (2008) Agricultural expansion within Kimanaelectric fences and implications for natural resource conservation aroundAmboseli National Park, Kenya. Journal of Arid Environments 72: 2179–2192.

50. Okello MM (2009) Contraction of wildlife dispersal area and displacement by human activities in Kimana Group Ranch near Amboseli National Park, Kenya. Open Conservation Biology Journal 3: 49–56.

51. Reid RS, Kaelo D, Galvin KA, Harmon R (2016) Pastoral Wildlife Conservancies in Kenya: A Bottom-up Revolution in Conservation, Balancing Livelihoods and Conservation? Proceedings of the International Rangelands Congress 18-22 July 2016, Saskatoon, Canada.

52. Lamprey RH, Reid RS (2004) Expansion of human settlement in Kenya’s Masai Mara: what future for pastoralism and wildlife?. Journal of Biogeography 31:997–1032.

53. Ogutu JO, Piepho HP, Reid RS, Rainy ME, Kruska RL, Worden JS,Hobbs NT (2010) Large herbivore responses to water and settlements in savannas. Ecological Monographs 80: 241–266.

54. Ogutu JO, Reid RS, Piepho H-P, Hobbs NT, Rainy ME, Kruska RL, Nyabenge M (2014). Large herbivore responses to surface water and land use in an East African savanna: implications for conservation and human-wildlife conflicts. Biodiversity and Conservation 23: 573–596.

55. Jolly GM (1969) Sampling methods for aerial censuses of wildlife populations. East African Agricultural and Forestry Journal 34(sup1): 46–49.

56. Burnham KP, Anderson DR (2002) Model selection and multimodel inference: a practical information-theoretic approach. Springer Science and Business Media.

57. SAS Institute Inc (2018) SAS system for windows. Cary, North Carolina, USA.

58. Stelfox JG, Peden DG, Epp H, Hudson RJ, Mbugua SW, Agastiva JL, Amuyunzu C L (1986) Herbivore Dynamics in Southern Narok, Kenya. Journal of Wildlife Management 50: 339–347.

59. Maddock L (1979) The “migration” and grazing succession. Pp. 104–129 In: ARE Sinclair, M Norton-Griffiths (1979), University of Chicago Press, Chicago, USA.

60. Hopcraft JGC, Morales J, Beyer H, Borner M, Mwangomo E, Sinclair ARE, Olff H, Haydon D (2014) Competition, predation, and migration: individual choice patterns of Serengeti migrants captured by hierarchical models. Ecological Monographs 84: 355–372.

61. Sinclair ARE (1973) Population increases of buffalo and wildebeest in the Serengeti. African Journal of Ecology 11: 93–107.

62. Sinclair ARE, Norton-Griffiths M (1982) Does competition or facilitation regulate ungulate populations in the Serengeti? A test of hypotheses. Oecologia. 53: 364–369.

63. Sidney J (1965) The past and present distribution of some African ungulates. Transactions of the Zoological Society of London 30: 1–396.

64. Jenkins P (2001) Wildlife use in World War II. In: An impossible dream, some of Kenya’s last colonial wardens recall the game department in the closing years of the British Empire, Pages 39–41, eds, I Parker and S Bleazard. Librario Publishing Limited.

65. Talbot LM, Stewart DRM (1964) First wildlife census of the entire Serengeti-Mara Region, East Africa. Journal of Wildlife Management 28: 815–827.

66. Talbot LM, Talbot MH (1961) Preliminary observations on the population dynamics of wildebeest in Narok District, Kenya. East African Agricultural and Forestry Journal 27: 108–116.

67. Simon N (1962) Between the sunlight and the Thunder: The Wild Life of Kenya. London: Collins.

68. Serneels S, Lambin EF (2001) Impacts of Land-use changes on the wildebeest migration in the northern part of the Serengeti-Mara ecosystem. Journal of Biogeography 28: 391–407.

69. Thomson J (1885) Through Masailand. Samson Low, Marston, Searle and Rivington, London.

70. Game Report (GAR) (1907) Game Annual Report 1906-07. Kenya National Archives.

71. Game Report (GAR) (1910) Game Report and lists of game killed 1909-1910. Kenya National Archives.

72. Heller E (1913) The white rhinoceros. Smithsonian Miscellaneous Collections 61: 1–77.

73. Roosevelt T, Heller E (1914) Life Histories of African Game Animals. Charles Scribner’s Sons, New York.

74. Meinertzhagen R (1957) Kenya Diary (1902-1906). Edinbugh: Oliver and Boyd.

75. Stewart DRM, Stewart J (1963) The distribution of some large mammals in Kenya. Journal of East African Natural History Society 24:1–52.

76. Percival AB (1928) A game Ranger on Safari. Nesbit Co Ltd, London.

77. Ogutu JO, Kuloba B, Piepho H-P, Kanga E (2017) Wildlife population dynamics in human-dominated landscapes under community-based conservation: Example of Nakuru Wildlife Conservancy, Kenya. PLos One 12(1), e0169730.

78. Estes RD, Atwood JL, Estes AB (2006) Downward trends in Ngorongoro Crater ungulate populations 1986–2005: Conservation concerns and the need for ecological research. Biological Conservation 131: 106–120.

79. Oates L, Rees PA (2013) The historical ecology of the large mammal populations of Ngorongoro Crater, Tanzania, east Africa. Mammal Review 43: 124–141.

80. Foster JB, Kearney D (1967) Nairobi National Park game census, 1966. East African Wildlife Journal 5: 112–120.

81. Foster JB, Coe MJ (1968) The biomass of game animals in Nairobi National Park, 1960-66. Journal of Zoology 155: 413–25.

82. Gichohi HW (1996) The Ecology of a truncated ecosystem, The Athi-Kapiti Plains. PhD Thesis, University of Leicester, Leicester.

83. Gichohi H (2000) Functional relationships between parks and agricultural areas in East Africa: the case of Nairobi National Park. In: Prins HHT, Grootenhuis JG, Thomas TD (Eds.) Wildlife Conservation by Sustainable Use.

84. De Beaton KP (1949) A warden’s diary. East African Standard.

85. Percival AB (1924) A Game Ranger’s Note Book. Nisbet, London.

86. Hillman JC, Hillman AK (1977) Mortality of wildlife in Nairobi National Park, during the drought of 1973–1974. African Journal of Ecology 15: 1–18.

87. Simon N (2001) New Directions in the 1950s. In: An impossible dream, some of Kenya’s last colonial wardens recall the game department in the closing years of the British Empire, Pages 83–92, eds, I Parker and S Bleazard. Librario Publishing Limited. Kinloss, Scotland.

88. Stanley J (2000) The Machakos Wildlife Forum: The story from a woman on the land. In Wildlife Conservation by Sustainable Use (pp. 13–20). Springer Netherlands.

89. Stabach JA (2015) Movement, resource selection, and the physiological stress response of white-bearded wildebeest. PhD Thesis, Colorado State University, Fort Collins, USA.

90. Sinclair ARE, Dublin H, Borner M (1985) Population regulation of Serengeti Wildebeest: a test of the food hypothesis. Oecologia 65: 266–268.

91. Mduma SAR, Sinclair ARE, Hilborn R (1999) Food regulates theSerengeti wildebeest: a 40-year record. Journal of Animal Ecology 68: 1101–1122.

92. Hobbs NT, Galvin KA, Stokes CJ, Lackett JM, Ash AJ, Boone RB, Reid RS, Thornton PK (2008) Fragmentation of rangelands: Implications for humans, animals, and landscapes. Global Environmental Change 18:776–785.

93. Western D (2007) A half a century of habitat change in Amboseli National Park, Kenya. African Journal of Ecology 45: 302–310.

94. Western D (2007) The ecology and changes of the Amboseli ecosystem. Recommendations for planning and conservation. Amboseli Conservation Program Report, 53pp. ACC, Nairobi, Kenya.

95. Mose VN, Nguyen-Huu T, Western D, Auger P, Nyandwi C (2013) Modelling the dynamics of migrations for large herbivore populations in the Amboseli National Park, Kenya. Ecological Modelling 254: 43–49.

96. Andere DK (1981) Wildebeest Connochaetes taurinus (Burchell) and its food supply in Amboseli Basin. African Journal of Ecology 19: 239–250.

97. Stabach JA, Wittemyer G, Boone RB, Reid RS, Worden JS (2016) Variation in habitat selection by white-bearded wildebeest across different degrees of human disturbance. Ecosphere 7(8):e01428. 10.1002/ecs2.1428.

98. Bhola N, Ogutu JO, Said MY, Olff H (2012) Herbivore hotspots in the Mara Region of Kenya in relation to land use. Journal of Animal Ecology 81: 1268–1287.

99. Morara MK, MacOpiyo L, Kogi-Makau W (2014) Land use, land cover change in urban pastoral interface. A case of Kajiado County, Kenya. Journal of Geography and Regional Planning 7: 192–202.

100. Reid RS, Rainy M, Ogutu JO, Kruska RL, Kimani K, Nyabenge M, McCartney M, Kshatriya M, Worden J, Ng’ang’a L, Owuor J, Kinoti J, Njuguna E, Wilson CJ, Lamprey R (2003) People, wildlife and livestock in the Mara ecosystem: The Mara Count 2002. International Livestock Research Institute, Nairobi, Kenya. Available online at: https://www.researchgate.net/profile/Joseph_Ogutu2/publication/266852551_People_wildlife_and_livestock_in_the_Mara_Ecosystem_the_Mara_Count_2002/links/54508a6d0cf249aa53da977c.pdf

101. Løvschal M, Bøcher PK, Pilgaard J, Amoke I, Odingo A, Thuo A, Svenning JC (2017) Fencing bodes a rapid collapse of the unique Greater Mara ecosystem. Nature Scientific Reports 7:41450. DOI: 10.1038/srep41450.

102. Williamson D, Williamson J (1985) Botswana’s fences and the depletion of Kalahari wildlife. Oryx 18: 218–222.

103. Tambling CJ, Du Toit JT (2005) Modelling wildebeest population dynamics: implications of predation and harvesting in a closed system. Journal of Applied Ecology 42: 431–441.

104. Western D (2010) The Worst Drought: Tipping point or Turning point. Swara 2:16–20.

105. Jones KE, Patel NG, Levy MA, Storeygard A, Balk D, Gittleman JL, Daszak P (2008) Global trends in emerging infectious diseases. Nature 451: 990–993.

106. Mills JN, Gage KL, Khan AS (2010) Potential influence of climate change on vector-borne and zoonotic diseases: a review and proposed research plan. Environmental Health Perspectives 118:1507–1514.

107. Bryony AJ, Grace D, Kock R, Alonso S, Rushton J, Said MY, McKeever D, Mutua F, Young J, McDermott J, Pfeiffer DO (2013) Zoonosis emergence linked to agricultural intensification and environmental change. Proceedings of the National Academy of Science 110(21): 8399–8404. http://dx.doi.org/10.1073/pnas.1208059110

108. Mduma SAR, Hilborn R, Sinclair ARE (1998) Limits to exploitation of Serengeti wildebeest and implications for its management. Pp 243–265 in Dynamics of tropical communities. Newbury DM, Prins, HHT Brown N (Eds). Oxford: Blackwell Science.

109. Rentsch D, Packer C (2015) The effect of bushmeat consumption on migratory wildlife in the Serengeti ecosystem, Tanzania. Oryx 49: 287–294.

110. Knapp EJ (2012) Why poaching pays: a summary of risks and benefits illegal hunters face in Western Serengeti, Tanzania. Tropical Conservation Science 5: 434–445.

111. Msoffe FU, Ogutu JO, Kaaya J, Bedelian C, Said MY, Kifugo SC, Reid RS, Neselle M, van Gardingen P, Thirgood S (2010) Paricipatory wildlife surveys in communal lands: a case study from Simanjiro, Tanzania. African Journal of Ecology 48: 727–735.

112. Lee DE, Kissui BM, Kiwango YA, Bond ML (2016) Migratory herds of wildebeests and zebras indirectly affect calf survival of giraffes. Ecology and Evolution 6: 8402–8411.

113. Bedelian C, Ogutu JO (2017) Trade-offs for climate-resilient pastorallivelihoods in wildlife conservancies in the Mara ecosystem, Kenya. Pastoralism 7: 10.

114. Norton-Griffiths M, Said MY (2010) The future for wildlife on Kenya’s rangelands: an economic perspective. Wild Rangelands: Conserving wildlife while mainting livestock in semi-arid ecosystems, p. 367–392.

115. KWS 2010 Aerial total count: Amboseli – West Kilimanjaro Natron cross border landscape, Wet season, March 2010. Available at https://www.kws.go.ke/kws/sites/default/files/Amboseli%20West%20Kilimanjaro%20and%20Magadi%20-%20Natron%20Cross%20Border%20Landscape%20March%202010%20Wet%20Season.pdf.

116. Thompson DM, Homewood K (2002) Entrepreneurs, elites and exclusion in Masailand: trends in wildlife conservation and pastoral development. Human Ecology 30: 107–138.

117. McCabe JT, Leslie PW, DeLuca L (2010) Adopting cultivation to remain pastoralists: The diversification of Masai livelihoods in northern Tanzania. Human Ecology 38: 321–334.

118. Boone RB, Galvin KA, Thornton PK, Swift DM, Coughenour MB (2006) Cultivation and conservation in Ngorongoro conservation area, Tanzania. Human Ecology 34: 809–828.

119. Norton-Griffiths M, Said MY, Serneels S, Kaelo DS, Coughenour M, Lamprey RH, Thompson DM, Reid RS (2008) Land use economics in the Mara Area of the Serengeti Ecosystem. Pp. 379–416 In: Serengeti III: The future of an ecosystem, Eds ARE Sinclair, C Packer, SAR Mduma, JM Fryxell. University of Chicago Press.

120. Mundia NC, Murayama Y (2009) Analysis of land use/cover changes and animal population dynamics in a wildlife sanctuary in East Africa. Remote Sensing 1: 952–970.

121. Norton-Griffiths M (1996) Property rights and the marginal wildebeest: An economic analysis of wildlife conservation options in Kenya. Biodiversity and Conservation 5: 1557–1577.

122. Homewood K (2009) Policy and practice in Kenya rangelands: Impacts on livelihoods and wildlife. Pages 335–367 in K Homewood, P Kristjanson, PC Trench, editors. Staying Masai? Livelihoods, conservation and development in East African rangelands. Springer, New York.

123. Western D, Groom R, Worden J (2009) The impact of subdivision and sedentarization of pastoral lands on wildlife in an African savanna ecosystem. Biological Conservation 142: 2538–2546.

124. Kideghesho JR (2002) Trends in areas adjacent to Tarangire National Park, Tanzania: What Community-Based land use planning can offer? Kakakuona[Jan-March]. Ministry of Natural Resources and Tourism, Tanzania.

125. Osano PM, Said MY, Leeuw J, Ndiwa N, Kaelo D, Schomers S, Ogutu JO (2013) Why keep lions instead of livestock? Assessing wildlife tourism‐based payment for ecosystem services involving herders in the Masai Mara, Kenya. Natural Resources Forum 37: 242–256.

126. Blackburn S, Hopcraft JGC, Ogutu JO, Matthiopoulos J, Frank L (2016) Human–wildlife conflict, benefit sharing and the survival of lions inpastoralist community‐based conservancies. Journal of Applied Ecology 53: 1195–1205.

127. Western D, Russell S, Cuthill I (2009) The status of wildlife in protected areas compared to non-protected areas of Kenya. PloS One 4: e6140.

128. Dougherty LS (2014) The ecological viability of wildlife conservancies in the Mara ecosystem. Transfer report to spatial ecology and land use unit, faculty of health and life sciences, Oxford Brookes University. Unpublished Report.

129. Nkedianye D, Radeny M, Kristjanson P, Herrero M (2009) Assessing returns to land and changing livelihood strategies in Kitengela Pages 115–150 in K Homewood, P Trench, and P Kristjanson, editors. Staying Masai? Livelihoods, Conservation and Development in East African Rangelands. Springer-Verlag, London.

130. de Leeuw JM, Said MY, Kifugo S, Ogutu JO, Osano P, de Leeuw J (2014) Spatial variation in the willingness to accept payments for conservation of a migratory wildlife corridor in the Athi-Kaputiei Plains, Kenya. Ecosystem Services 8: 16–24.

131. Matiko D (2014) Wildlife conservation leases are considerable conservation options outside Protected Areas: The Kitengela-Nairobi National Park Wildlife Conservation Lease Program. Journal of Ecosystem and Ecography. 4:2. http://dx.doi.org/10.4172/2157-7625.1000146.

132. Nelson F, Foley C, Foley LS, Leposo A, Loure E, Peterson D, Peterson MPeterson T, Sachedina H,Williams A (2010) Payments for ecosystemservices as a framework for community-based conservation in NorthernTanzania. Conservation Biology 24: 78–85.

133. Morrison TA, Bolger DT (2014) Connectivity and bottlenecks in a migratory wildebeest Connochaetes taurinus population. Oryx 48: 613–621.

134. Norton-Griffiths, M (2000) Wildlife losses in Kenya: An analysis of conservation policy. Natural Resources Modelling 13: 13–34.

135. Republic of Kenya (2013) The Wildlife Conservation and Management Act, 2013. Kenya Gazette Supplement No. 181, Acts No. 47, Sixth Schedule. Nairobi. Available at: http://kenyalaw.org/kl/fileadmin/pdfdownloads/Acts/WildlifeConservationandManagement%20Act2013.pdf. Kenya Gazette Supplement No. 18/ (Acts No. 47). Republic of Kenya.

136. Ministry of Tourism and Wildlife (2018) National Wildlife Strategy 2030. Govenrment of Kenya Publication, Nairobi.

137. Ojwang’ GO, Wargute PW, Said MY, Worden JS, Davidson Z, Muruthi P,Kanga E, Ihwagi F, Okita-Ouma B (2017) Wildlife migratory corridors and dispersal areas: Kenya rangelands and coastal terrestrial ecosystems. Government of the Republic of Kenya, Nairobi.

138. United Republic of Tanzania (2018) Wildlfie Conservation (Wildlife corridors, dispersal areas, buffer zones and migratory routes) Regulations 2018. Government Printer, Dar es Saalam.

139. Ogutu J O, Owen-Smith N, Piepho H P, Kuloba B, Edebe J (2012) Dynamics of ungulates in relation to climatic and land use changes in an insularized African savanna ecosystem. Biodiversity Conservation 21:1033–1053.

140. Swynnerton GH (1958) Fauna of the Serengeti National Park. Mammalia 22: 435–450.

141. Grizmek B, Grizmek M (1960) Serengeti shall not die. Hamish Hamilton, Ltd. London. 344pp.

142. Stewart D RM, Talbot LM (1962) Census of wildlife in the Serengeti, Mara and Loita Plains. East African Agricultural and Forestry Journal 28: 58–60.

143. Anderson GD, Talbot, LM (1965) Soil factors affecting the distribution of the grassland types and their utilisation by wild animals on the Serengeti plains, Tanganyika. Journal of Ecology 53: 33–56.

144. Watson RM (1967) The population ecology of the wildebeest (Connochaetes taurinus albojubatus Thomas) in the Serengeti. PhD Thesis, Cambridge University.

145. Bell R V H (1971) A grazing ecosystem in the Serengeti. Scientific American 224: 86–93.

146. Kreulen D (1975) Wildebeest habitat selection on the Serengeti Plains, Tanzania, in relation to Calcium and lactation. African Journal of Ecology 13: 297–304.

147. McNaughton S J (1976) Serengeti migratory wildebeest: Facilitation of energy flow by grazing. Science 191 (4222): 92–94.

148. Hilborn R, ARE Sinclair (1979) A simulation of the wildebeest population, other ungulates and their predators. Pages 287–309 in ARE Sinclair and MNorton-Griffith, editors. Serengeti: dynamics of an ecosystem. University ofChicago Press, Chicago.

149. Sinclair ARE (1979) The eruption of the ruminants. Pp 82–103 In: A R E Sinclair and M Norton-Griffith, editors. Serengeti: dynamics of an ecosystem. University of Chicago Press, Chicago.

150. Sinclair ARE, Norton-Griffiths M (1979) Serengeti: Dynamics of an ecosystem. University of Chicago Press, Chicago, USA.

151. Bell RVH (1982) The effect of soil nutrient availability on community structure in African ecosystems. In: Ecology of tropical savannahs, p. 193–216. ed. BJ Huntley, Walker BH. Springer, New York.

152. Broten MD, Said MY (1995) Population trends of ungulates in and around Kenya’s Masai Mara Reserve. In: Serengeti II: Dynamics, Management, andConservation of an Ecosystem, p. 169–193. ed. ARE Sinclair. P Arcese.Univ. of Chicago Press, Chicago.

153. Fryxell JM (1995) Aggregation and migration by grazing ungulates in relation toresources and predators. In: Serengeti II. Dynamics, Management, andConservation of an Ecosystem, p. 257–273. ed. ARE Sinclair, P Arcese. Univ. of Chicago Press, Chicago.

154. Murray MG (1995) Specific nutrients requirements and migration of wildebeest. Pp 231–56 In: Serengeti II; dynamics, management and conservation of anecosystem. University of Chicago Press, Chicago, USA.

155. Wilmshurst JF, Fryxell JM, Farm BP, Sinclair ARE, Henschel CP (1999) Spatial distribution of Serengeti wildebeest in relation to resources. Canadian Journal of Zoology 77: 1223–1232.

156. Gereta E, Wolanski E, Chiombola EAT (2003) Assessment of the environmental, social and economic impacts on the Serengeti ecosystem of the developments in the Mara. River catchment in Kenya. Amala Project Report, 59pp. TANAPA, FZS, Arusha, Tanzania.

157. Musiega DE, Kazadi SN (2004) Simulating the East African wildebeestmigration patterns using GIS and remote sensing. African Journal of Ecology 42: 355–62.

158. Boone RB, Thirgood SJ, Hopcraft JGC (2006) Serengeti wildebeest migratory patterns modeled from rainfall and new vegetation growth. Ecology 87: 1987–94.

159. Sinclair ARE, Mduma SA, Hopcraft JGC, Fryxell J M, Hilborn R A Y, Thirgood S (2007) Long-Term Ecosystem Dynamics in the Serengeti: Lessons for Conservation. Conservation Biology 21: 580–590.

160. Holdo R M, Holt R D, Fryxell J M (2009) Grazers, browsers, and fire influence the extent and spatial pattern of tree cover in the Serengeti. Ecological Applications 19: 95–109.

161. Bhola N, Ogutu J O, Piepho H-P, Said MY, Reid RS, Hobbs NT, OlffH (2012b) Comparative changes in density and demography of largeherbivores in the Masai Mara Reserve and its surrounding human-pastoralranches in Kenya. Biodiversity Conservation 21: 1509–1530.

162. Bedelian C (2014) Saving the Great Migrations: Declining wildebeest in East Africa? Environmental Development 9: 101–109.

163. Stabach JA, Boone RB, Worden JS, Florant G (2015) Habitat disturbanceeffects on the physiological stress response in resident Kenyan white-bearded wildebeest (Connochaetes taurinus). Biological Conservation 182:177–186.

164. Ottichilo WK (2000) Wildlife Dynamics: An Analysis of Change in the Masai Mara Ecosystem of Kenya. PhD Dissertation, ITC, The Netherlands.

165. Sheehan MM (2016) Determining drivers for wildebeest (Connochaetes taurinus) distribution in the Masai Mara National Reserve and surrounding Group Ranches (Doctoral dissertation, Miami University).

166. McCutcheon JT (1910) In Africa. The Bobbs-Merrill Company. Indianapolis, USA.

167. Foster JB, McLaughlin R (1968) Nairobi National Park game census, 1967. EastAfrican Wildlife Journal 6: 152–54.

168. Casebeer RL, Koss GG (1970) Food habits of wildebeest, zebra, hartebeestand cattle in Kenya Masailand African Journal of Ecology 8: 25–36.

169. Petersen JCB, Casebeer RL (1972) Distribution, population status and group composition of wildebeest (Connochaetes taurinus Burchell) and zebra (Equus burchelli Gray) on the Athi-Kapiti plains, Kenya. Wildlife Management Project. UNDP/FAO KEN/71 /526, Project Working Document No. 1.

170. Casebeer RL, Mbai HJ (1974) Animai mortality 1973/74. Kajiado District, FAOProject DP/KEN/71/526. Working Document. Report No. 5.

171. Owaga M L (1975) The feeding ecology of wildebeest and zebra in Athi-Kaputei Plains. African Journal of Ecology 13: 375–83.

172. Hillman JC (1979) The biology of the eland (Taurotraaus oryx Pallas) in the wild. PhD. University of Nairobi.

173. Trzebinski E (1985) The Kenya Pioneers. Cox and Wyman Ltd, Great Britain.

174. Gichohi HW (2003) Direct payments as a mechanism for conserving important wildlife corridor links between Nairobi National Park and its wider ecosystem: The Wildlife Conservation Lease Program. In Vth World Parks Congress.

175. Ego WK, Mbuvi D M, Kibet PFK (2003) Dietary composition ofwildebeest (Connochaetes taurinus), kongoni (Alcephalus buselaphus) andcattle (Bos indicus), grazing on a common ranch in south‐central Kenya. African Journal of Ecology 41: 83–92.

176. Imbahale SS, Githaiga J M, Chira RM, Said MY (2008) Resource utilization by large migratory herbivores of the Athi-Kapiti ecosystem. African Journal of Ecology 46: 43–51.

177. Croze H (1978) Aerial surveys undertaken by the Kenya Wildlife Management Project: meththodoiogies and results. FAO Project DP/KEN/71/526. Working document. Report No. 16.

178. Campbell DJ, Gichohi H, Mwangi A, Chege L (2000) Land Use Conflicts in Kajiado District, Kenya. Land Use Policy 17: 337–48.

179. Worden J, Reid RS, Gichohi H (2003) Land-use impacts on large wildlife and livestock in the swamps of the Greater Amboseli Ecosystem, Kajiado District, Kenya Lucid Project. International Livestock Research Institute, Nairobi, Kenya. https://cgspace.cgiar.org/bitstream/handle/10568/1901/Lucid_wp27_part1.pdf?sequence=1.

180. Okello M M (2005) Land use changes and human–wildlife conflicts in theAmboseli Area, Kenya. Human Dimensions of Wildlife 10: 19–28.

181. Sitati N, Lekishon K, Bakari S, Warinwa F, Mwiu S N, Gichohi N, Mukeka J (2014). Wildebeest (Connochaetes taurinus) Population densities and distribution in dry and wet season in the Kilimanjaro landscape. Natural Resources 5: 810.

182. Galanti V, Tosi G, Rossi R, Foley C (2000) The Use of GPS radio-collars to track elephants (Loxodonta africana) in the Tarangire National Park (Tanzania). Hystrix 11: 27–37.

183. TAWIRI (2001) Tarangire Ecosystem: Wet Season Systemmatic ReconnaisanceFlight Count, May 2001. TAWIRI, Arusha, Tanzania.

184. Gereta E, Meing’ataki GEO, Mduma SAR, Wolanski E (2004) The role ofwetlands in wildlife migration in the Tarangire ecosystem, Tanzania. Wetlands Ecology and Management 12: 285–99.

185. Newmark WD (2008) Isolation of African protected areas. Frontiers in Ecology and the Environment 6:321–8.

